# Crop-OCT: a Fully Integrated Imageomics Pipeline to Identify Regional and Focal Retinopathy in Murine Models

**DOI:** 10.64898/2026.02.27.708333

**Authors:** Danielle R. Little, Abbas Shirinifard, Marybeth Lupo, Chih-Hsuan Wu, Haoran Chen, Mellisa R. Clemons, Michael Maclean, Olivia J. Marola, Gareth R. Howell, Cai Li, Michael A. Dyer

**Affiliations:** Departments of Developmental Neurobiology, St. Jude Children’s Research Hospital, Memphis, TN, USA; Biostatistics, St. Jude Children’s Research Hospital, Memphis, TN, USA; The Jackson Laboratory, Bar Harbor, ME, USA

## Abstract

Imageomics uses machine learning to accelerate our understanding of biological traits and human disease processes. Some of the earliest imageomics applications used deep learning to assess human diseases. For example, retinal fundus images were analyzed to diagnose diabetic retinopathy. The imaging modality optical coherence tomography (OCT) is widely used to diagnose and monitor the progression of retinopathy in patients and preclinical models. The standardized instrumentation and image format of OCT lends itself to imageomics, but generalizable, automated pipelines for segmentation and quantitation of large numbers of OCT images are still in early development. Here, we present the automated, end-to-end pipeline Crop-OCT that extracts features from thousands of OCT images, while preserving their location within the eye. We used the Crop-OCT pipeline on a diverse dataset, including 13 genetic models of retinopathy, with more than 20,000 OCT images, which allowed us to analyze nearly 6 million measured features. The pipeline was generalized on an independent dataset that was analyzed in a blinded manner. The pipeline enabled us to monitor ocular changes associated with aging and progression of diseases, such as retinitis pigmentosa, Leber congenital amaurosis, achromatopsia, Stargardt disease, diabetic retinopathy, and age-related macular degeneration. We also characterized heterogeneity across animals and identified regional and focal lesions. Our pipeline will unify feature extraction for preclinical models of retinal disease and serve as a foundation for future multimodal data integration for artificial intelligence applications based on imageomics.

## INTRODUCTION

The emerging field of imageomics extracts quantitative, biological information from images at scale, including microscopic, medical, and species-tagged natural world images. Investigators have developed artificial intelligence (AI) methods to extract features from thousands of hematoxylin and eosin (H&E)-stained whole-slide microscopy images for digital pathology(1, 2). In the fields of ecology and biodiversity, AI tools are being developed to fill major knowledge gaps, including automated identification of previously unidentified taxa and dynamic biogeographic distribution of species due to climate change(3). The most rapidly expanding area of imageomics is in diagnostic imaging–mediated diagnosis of cancer(4) and neurologic diseases(5, 6). One of the best-established applications of imageomics to human health is for diagnosis of retinopathy. The first FDA-approved autonomous AI tool for disease diagnosis is IDx-DR, which detects diabetic retinopathy on retinal fundus images(7, 8). In addition, ocular image biomarkers of diverse human diseases are being identified in large cohorts, such as the UK biobank(9–11).

Another common imaging tool used in ophthalmology is optical coherence tomography (OCT)(12, 13). This approach enables ophthalmologists to analyze ocular structures, diagnose retinopathy, and monitor disease progression(14). The advantage of OCT is that it is noninvasive and can quickly provide quantitative spatial data on the thickness of retinal layers and other ocular structures, such as the choroid and retinal pigment epithelium (RPE). OCT is widely used in preclinical models of ocular diseases, ranging from retinitis pigmentosa, to age-related macular degeneration (AMD), and diabetic retinopathy(15, 16). Applying imageomics to OCT data represents an important opportunity to distinguish various ocular diseases(17). Moreover, OCT is emerging as a foundation for oculomics—identifying ocular biomarkers for systemic diseases, including pulmonary hypertension, chronic kidney disease, and Alzheimer disease(9, 18).

For human OCT data, there are several large databases containing clinically annotated image datasets; however, there is no generalized model for segmentation of retinal layers, focal disruptions, or feature extraction for AI applications(19). For preclinical OCT data, most studies focus on individual murine models, and the segmentation and feature extraction may not be generalizable across diverse phenotypes(20–22). Aside from the quantitation of retinal layers, segmentation often focuses on disease-specific features(23). Development of an automated generalizable segmentation, quality control (QC), and feature-extraction pipeline will accelerate the application of AI tools to diagnose and monitor ocular disease and serve as a foundation for multimodal integration.

Here we developed an end-to-end pipeline for preclinical OCT segmentation, QC, feature extraction, and statistical analysis. We trained and tested our pipeline on 15 murine models of retinal disease over multiple time points (3–27 months of age). In total, 21,147 OCT images had 267 features extracted per image. We sampled 8 regions per eye, and the OCT images were spatially registered with the matched fundus images. We validated our results by manual annotation and histopathologic analysis in a blinded manner. From statistical analysis of the features extracted by our pipeline, we identified retina-wide changes and focal lesions. We demonstrated the generalized ability of our new pipeline on an external dataset from 2 unrelated murine models of heterogenous retinal degeneration.

## RESULTS

### Standardized OCT image acquisition across diverse models and time points

To develop an end-to-end pipeline for analyzing preclinical OCT images, we established large colonies of 13 genetic murine models of retinopathy and 2 inbred strains (**Table S1**). Using the Phoenix MICRON IV retinal imaging microscope, we acquired paired fundus and OCT images for each eye, across 3-27 months of age (**Figure 1A-D** and **Table S1**). Data were acquired from 4 mice (2 male, 2 female) per time point per strain for a total of 336 mice (**Table S1**). OCT images were acquired at 3 locations (superior, midline, inferior) across the eye to capture spatial heterogeneity for a total of 8873 OCT images (**Figure 1E**). Multiple OCT images were acquired at each location in each mouse, across all time points to overcome motion artifacts caused by breathing. We included models of retinitis pigmentosa, Leber congenital amaurosis, achromatopsia, Stargardt disease, diabetic retinopathy, and AMD (**Figure 1B** and **Table S1**).

**Figure 1.**
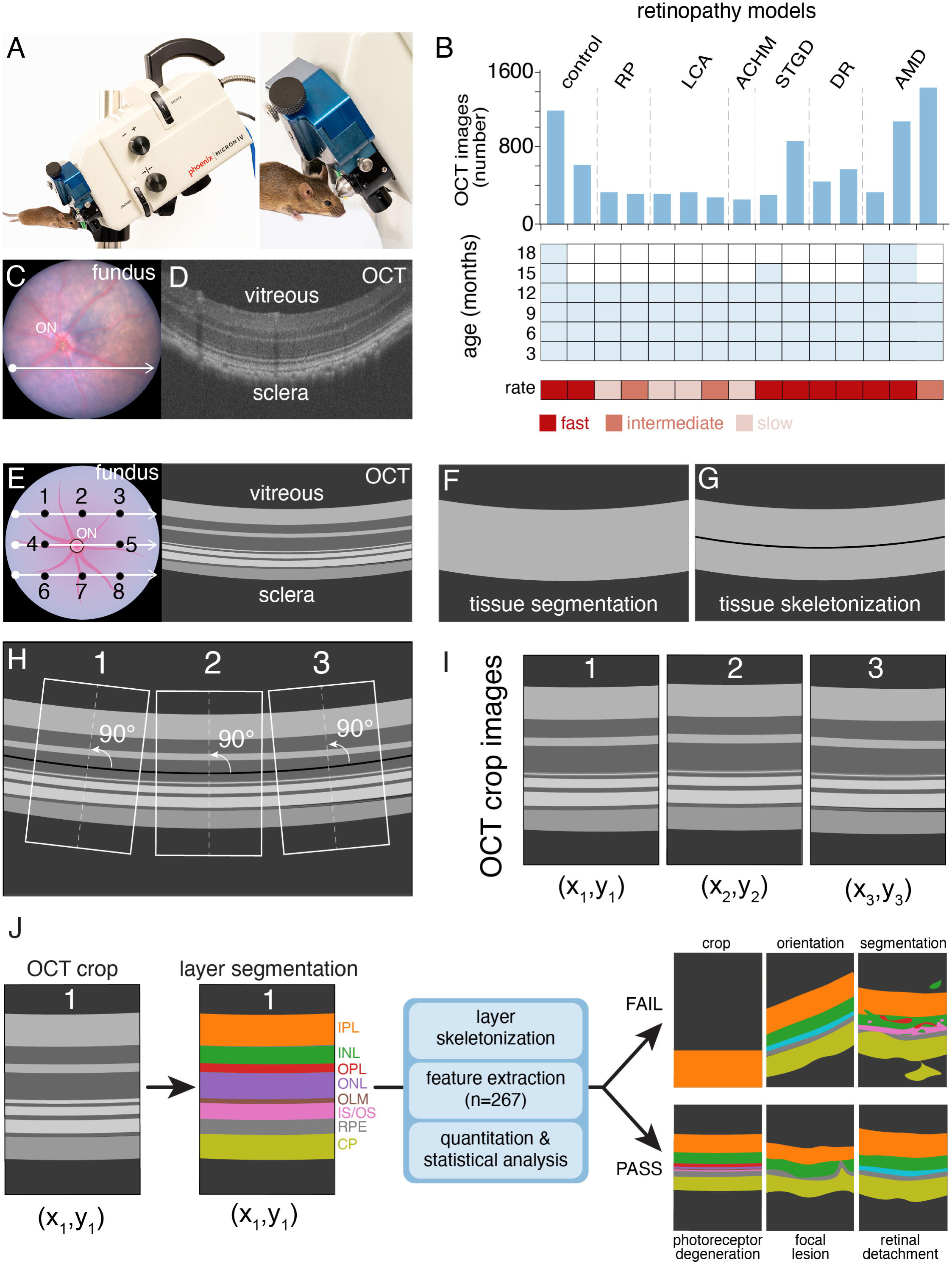
Development of an automated pipeline for OCT analysis. **A)** Photograph of the Phoenix MICRON IV retinal imaging microscope. **B)** Histogram of the number of OCT images for each of the 15 mouse strains in this study (upper panel). Heatmap of the time points at which OCT images were obtained (blue) for each strain (middle panel), and rate of disease progression (red) for each model (lower panel) are shown. **C,D)** Matched fundus (C) and OCT image (D) for a representative 6-month-old C57BL/6J mouse. **E)** Drawing depicting the 3 locations sampled in the superior, midline, and inferior positions for line scans (white arrows) and the respective locations where cropped OCT images (black dots) were sampled for further analysis. **F,G)** Drawings showing the Ilastik segmentation of tissue versus background (F) and tissue skeletonization (G). **H,I)** Drawings showing how the medial axis (white dashed line) perpendicular to the skeletonized line is used to reorient individual cropped OCT images, while preserving the *x*,*y*-coordinates. **J)** Diagram showing the segmentation of 8 individual ocular layers and the pipeline to skeletonize those layers, extract features, quantitate and analyze differences across images, eyes, and mice. Drawings of images that may fail or pass quality control from the C-OCT pipeline are shown. **Abbreviations:** ACHM, achromatopsia; AMD, age-related macular degeneration; CP, choroid plexus; DR, diabetic retinopathy; INL, inner nuclear layer; IPL, inner plexiform layer; IS/OS, inner segment/outer segment; LCA, Leber congenital amaurosis; OCT, optical coherence tomography; OLM, outer limiting membrane; ON, optic nerve; ONL, outer nuclear layer; OPL, outer plexiform layer; RP, retinitis pigmentosa; RPE, retinal pigment epithelium; STGD, Stargardt disease.

We trained an Ilastik model to segment the tissue (retina, RPE, and choroid) and defined the midline (**Figure 1F,G**)(24). We cropped 3 regions perpendicular to the midline for each OCT image to account for ocular curvature (**Figure 1H**). Therefore, each eye had 8 cropped images (excluding optic nerve head) with relative spatial position (*x*,*y*) preserved (**Figure 1E,I**). Next, we trained 2 Mamba-based models (Swin-UMamba models) by using our manual ground truth annotations to segment the 8 layers in our OCT-cropped images and to segment the optic nerve head in the fundus images (**Figure 1J**)(25). A total of 296 cropped OCT images and 219 fundus images from multiple genetic models and time points were annotated for training Swin-UMamba (**Figure 1E**, **Supplemental Information**). To validate the segmentation, we manually reviewed ∼300 cropped OCT images and fundus images per strain (**Supplemental Information**). All 22,384 cropped OCT images were segmented, and 267 features (**Table S2**) were extracted per image for a total of 5,976,261 data points. We used a subset of these features to set QC thresholds while retaining focal lesions associated with disease progression (**Figure 1J**, **Supplemental Information**). Overall, 94.5% (21,147/22,384) of the cropped OCT images passed our QC (**Figure S1**). There was no significant difference across the technical replicates for cropped OCT images (**Supplemental Information**), so we de-duplicated the data to minimize the impact of unequal technical replicates across individual eyes and mice. Finally, we performed statistical analysis of OCT features of individual eyes and mice, relative to age-matched controls at individual time points and with aging to monitor disease progression. This end-to-end pipeline is called Crop–optical coherence tomography (C-OCT).

### Quantitating Retinal Disease Progression from OCT Images

To analyze disease progression in our murine models of retinopathy, we evaluated the thickness of 8 retinal layers and compared the individual models to their age-matched genetic controls. For example, the *Rpe65^rd12/rd12^*strain is a spontaneous mutant model with a nonsense mutation that leads to photoreceptor cell death and blindness(26). At 3 months of age, the outer nuclear layer (ONL) was significantly thinner in the *Rpe65^rd12/rd12^* mice, relative to C57BL/6J animals, and it became progressively thinner over time (**Figure 2A-D**). As expected, the changes in the ONL were accompanied by progressive thinning of the inner and outer segments (IS/OS), the outer plexiform layer (OPL), and the RPE. These results were validated by histopathology images (**Figures 2C**, **S2**).

**Figure 2.**
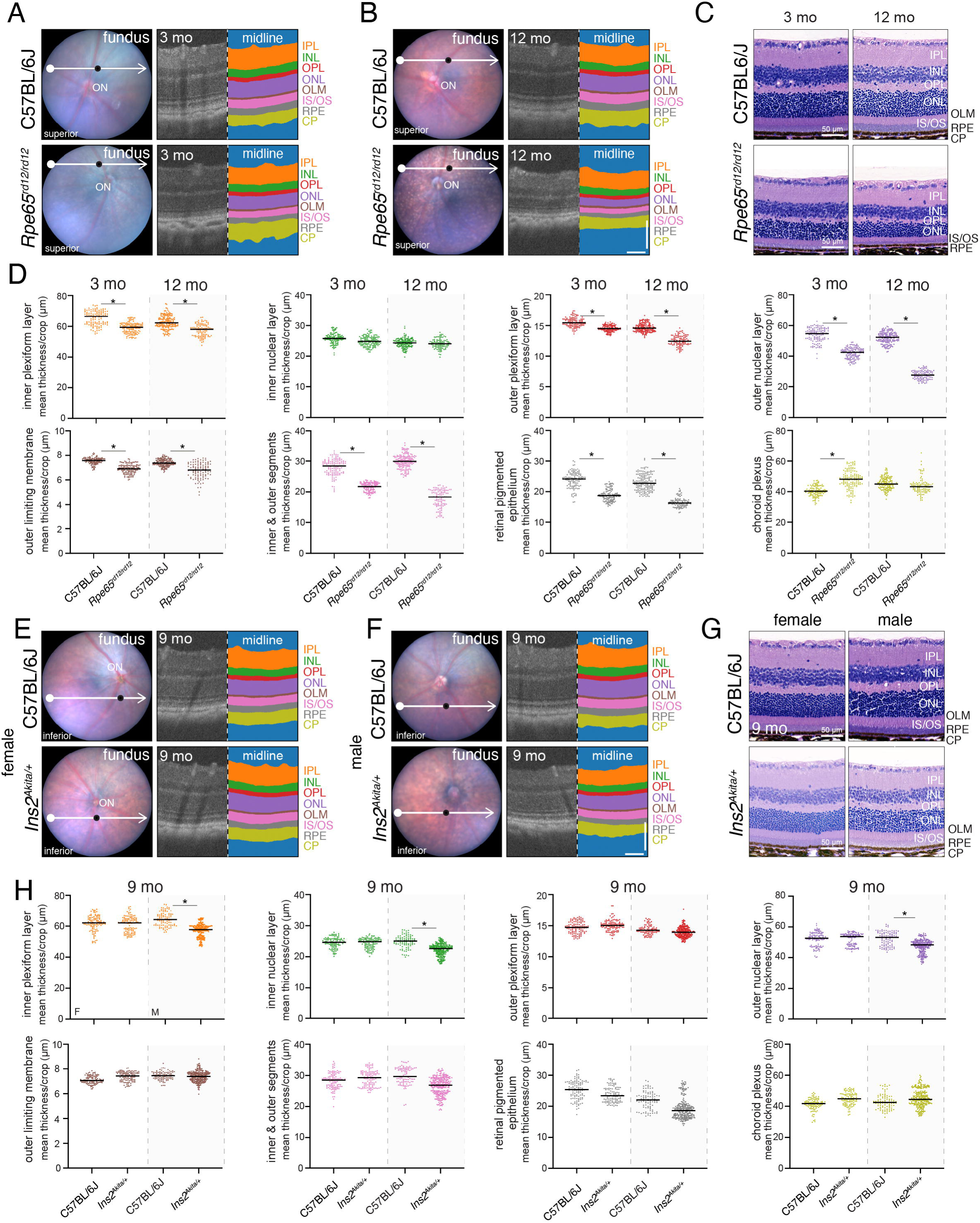
C-OCT can be used to monitor changes in ocular layer thickness with disease progression. A,B) Images of fundi with scan line (white) and paired midline OCT images (location: black dot on fundus images) with segmentation for a C57BL/6J mouse (upper panel) and a *Rpe65^rd12/rd12^*mouse (lower panel) as 3 months (A) and 12 months (B). **C)** H&E-stained histologic images of eyes from the mice shown in (A,B). **D)** Plots of average layer thicknesses for the strains shown in (A-C). Each dot is an individual cropped OCT image. Kruskal-Wallis tests with adjusted p-value, *p <0.05. **E,F)** Images representative of fundi with inferior scanlines (white arrows) and corresponding midline OCT image (location: black dot on fundi images) and segmentation of 9-month-old female C57BL/6J mice and female *Ins2^Akita/+^*mice (E). Images of corresponding male mice are shown in (F). **G)** H&E-stained histologic images of the mice shown in (E,F). **H)** Plots of average layer thicknesses for the strains shown in (E-G). Each dot is an individual OCT cropped image. Kruskal-Wallis tests with adjusted p-value, * p <0.05. **Abbreviations:** CP, choroid plexus; H&E, hematoxylin and eosin; INL, inner nuclear layer; IPL, inner plexiform layer; IS/OS, inner segment/outer segment; OLM, outer limiting membrane; ON, optic nerve head; ONL, outer nuclear layer; OPL, outer plexiform layer; RPE, retinal pigment epithelium. OCT image scale bars: 100 μm.

Our C-OCT pipeline also identified sex-influenced phenotypes. The *Ins2^Akita/+^* strain, which has a missense mutation in the *Ins2* gene, is a model for type 1 diabetes(27). Heterozygous male *Ins2^Akita/+^*mice have persistently elevated blood glucose, but female mice do not(27). The male mice develop diabetic retinopathy, which is accompanied by vascular leakage, ganglion cell loss, synaptic loss, and changes in photoreceptor function(28–30). In our quantitative assessment of the cropped OCT images, the male mice had significant thinning of the inner plexiform layer (IPL), which was consistent with ganglion cell loss and thinning of the IS/OS and ONL, causing photoreceptor dysfunction (**Figure 2E-H**). We also identified thinning of the RPE, which was consistent with vascular changes in the RPE and choroid associated with neovascularization.

These changes were confirmed in histologic images (**Figure 2G**). The data confirm that our C-OCT pipeline can be used to quantify both subtle and gross structural changes in ocular layers associated with disease progression from large imageomics datasets.

### Identifying Regional Heterogeneity in Retinopathy

To identify regional heterogeneity in ocular layer thickness, we preserved the spatial distribution of individual cropped OCT images (**Figure 1E,I**). The optic nerve head was automatically segmented in our C-OCT pipeline (**Figure S3A-C**), and the distance from the center of the segmented optic nerve head to the center of the cropped OCT image was linked to the individual image files (**Figure S3D**). The *x*,*y* coordinates for each cropped OCT image can be related to an anatomic location (**Figure S3D**). As described above, some images were eliminated because they did not pass our QC threshold, but for the majority (68%) of eyes, all 8 cropped images were retained (**Figure S3E,F**). We set a minimum threshold of 5 cropped images per eye with at least 2 *y*-locations for our C-OCT pipeline (**Figure S3E,F**). In total, 94.2% (618/654) of eyes met our threshold for analysis of regional heterogeneity.

We extracted 448 eye-level features for the 616 eyes and performed statistical analysis on the 275,968 data points to identify regional heterogeneity at each time point. In the *Tsc1^Lox/Lox^;Rho-iCre;OPN1LW-Cre* strain (*Tsc1*), a murine model of AMD, the *Tsc1* gene is conditionally inactivated in photoreceptors (**Table S2**)(31–34). This model showed the largest significant difference in standard error of the mean (SEM) for normalized ocular layer thickness, relative to age-matched controls and over time (**Figure 3A,B**). The ONL thickness was variable across mice at 12 months of age, with thinning in the superior region of the eye (**Figure 3C-E**). These results were validated at the individual eye level and in the histologic images (**Figure 3F-I**). Importantly, the *Tsc1* mice were not used in the original training for ocular layer and fundus segmentation, so this result demonstrates the generalized ability of the automated segmentation step of our C-OCT pipeline. The analysis of the *Tsc1* OCT images demonstrates the importance of preserving spatial information to identify phenotypes with regional heterogeneity.

**Figure 3.**
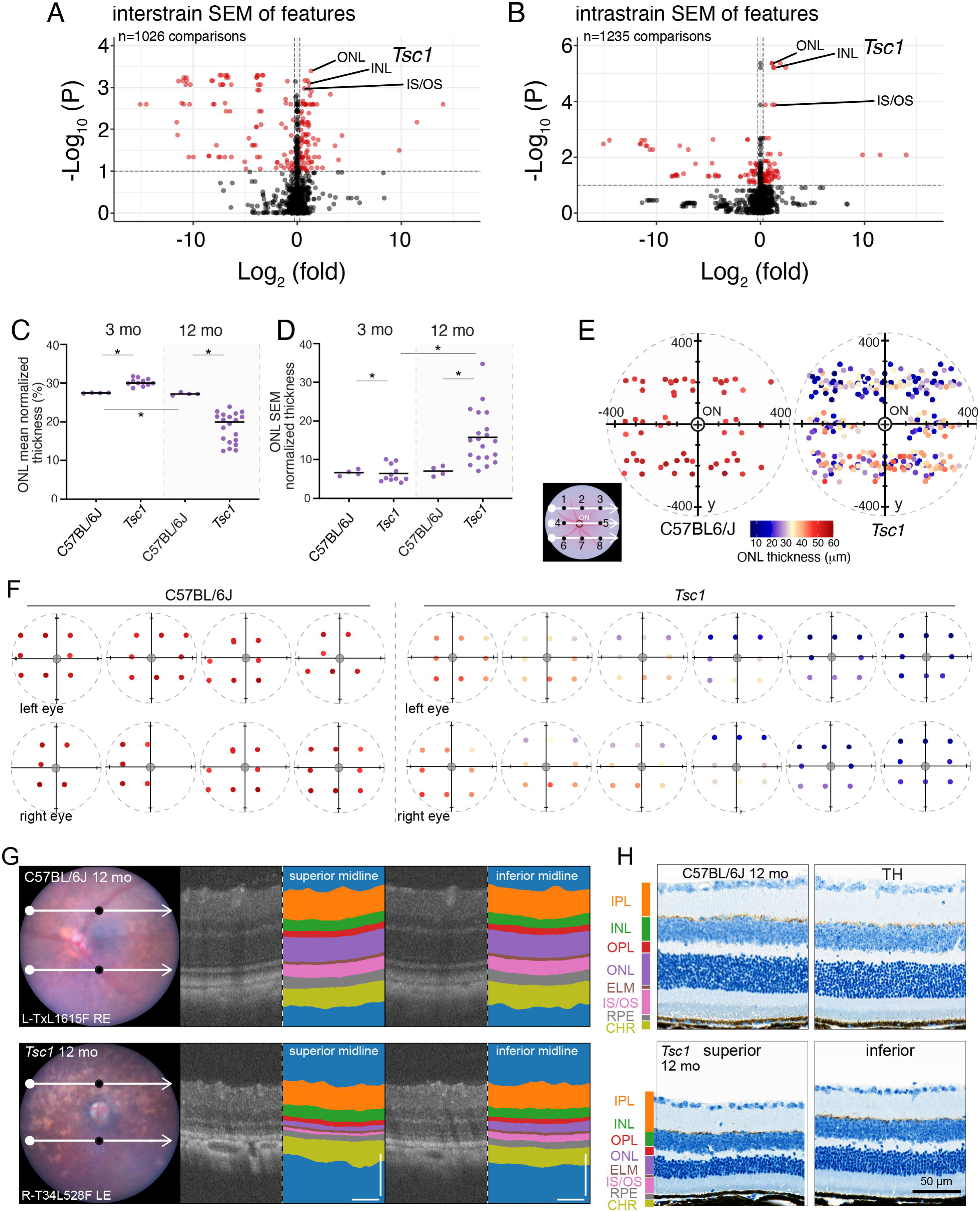
Identification of regional heterogeneity of ocular layer thickness by using C-OCT. **A)** Volcano plot of all C-OCT data standard error of the means (SEM) comparisons between mutant mice and their respective time point control mice with Kruskal-Wallis statistical test that was adjusted for multiple testing (p <0.01; red dots). Log_2_ (fold-change) was calculated between mutants and their respective genetic background–matched time point control mice. Highlighted are the SEM values for normalized average thickness of ONL, INL, and IS/OS for 12-month-old *Tsc1^lox/lox^;Rho*-iCre^Tg^;Tg-*OPN1MW*-Cre^Tg^ (n = 20 mice, 37 eyes) versus C57BL/6J (n = 4 mice, 8 eyes). **B)** Volcano plot of all C-OCT cohort comparisons within mutant mice compared across time within their respective genotypes with Kruskal-Wallis statistical test that was adjusted for multiple testing (p <0.01; red dots). Log_2_ (fold-change) was calculated between mutants and control mice at the 3-month time point. Highlighted are the SEM values for normalized average thickness of ONL, INL, and IS/OS in 12-month-old *Tsc1^lox/lox^;Rho*-iCre^Tg^;Tg-*OPN1MW*-Cre^Tg^ (n =20 mice, 37 eyes) versus 3-month-old C57BL/6J *Tsc1^lox/lox^;Rho*-iCre^Tg^;Tg-*OPN1MW*-Cre^Tg^ (n = 11 mice, 21 eyes). **C,D)** Plots of the ONL mean normalized thickness per mouse strain (C) and the ONL SEM of the normalized average thickness per mouse strain (D). Kruskal-Wallis adjusted *p-value <0.05. **E-F)** Eye map ONL thickness plot of all de-duplicated images (points) for each eye on the optic nerve coordinate grid, with the mean ONL thickness plotted by the hue of each point for all 12-month-old C57BL/6J (n = 4 mice, 8 eyes; left) and *Tsc1^lox/lox^;Rho*-iCre^Tg^;Tg-*OPN1MW*-Cre^Tg^ (n = 20 mice, 37 eyes; right). Individual mice eye maps are shown in (F). **G)** Images of representative fundi and paired superior midline (left) and inferior midline (right) OCT images of C57BL/6J (n = 4 mice, 8 eyes; upper panel) and *Tsc1^lox/lox^;Rho*-iCre^Tg^;Tg-*OPN1MW*-Cre^Tg^ (n = 20 mice, 37 eyes; lower panel). scale bar = 100 μm. **H)** Images of histologic sections used for immunohistochemical staining of 12-month-old C57BL/6J (n =4 mice; top panel) and *Tsc1^lox/lox^;Rho*-iCre^Tg^;Tg-*OPN1MW*-Cre^Tg^ (n = 4 mice; lower panel), superior (left) and inferior (right) OCT images. **Abbreviations:** INL, inner nuclear layer; IPL, inner plexiform layer; IS/OS, inner segment/outer segment; OLM, outer limiting membrane; ON, optic nerve head; ONL, outer nuclear layer; OPL, outer plexiform layer; RPE, retinal pigment epithelium; SEM, standard error of the mean; TH, tyrosine hydroxylase. OCT scale bars:100 μm.

### Identifying Focal Disruptions in Ocular Structures

To exploit the position information of individual cropped OCT images in our C-OCT pipeline, we developed methods to identify the most common types of focal disruptions. To identify retinal detachments, we added a segmentation mask to the pipeline (**Figure 4A** and **Supplemental Information**). Layer thickness was measured at each pixel across the image, and measurements were averaged, normalized, and used for statistical analysis of variation within images (**Figures 4B**, **S4**, and **Supplemental Information**). To further enhance our ability to identify local variation in ocular structures within individual OCT images, we measured the angle of the tangent to the medial axis (skeleton) of each layer in a sliding window across each OCT image (**Figure 4C**). Using these measurements for each image, we identified OCT images with focal disruptions in ocular structures (**Figures 4C**, **S4,** and **S5**). The most acute changes in the angle of the tangent of the medial axis were often associated with domain breaks and/or a bifurcation of one or more medial axis skeletons (**Figure 4D**). To identify focal lesions across our cohort, we performed unsupervised hierarchical clustering of a subset of normalized measurements from individual cropped OCT images (**Figure 4E, Table S3**). Those images with retinal detachments formed a separate cluster that included *Cep290^rd16/rd16^*mice (**Figure 4F**).

**Figure 4.**
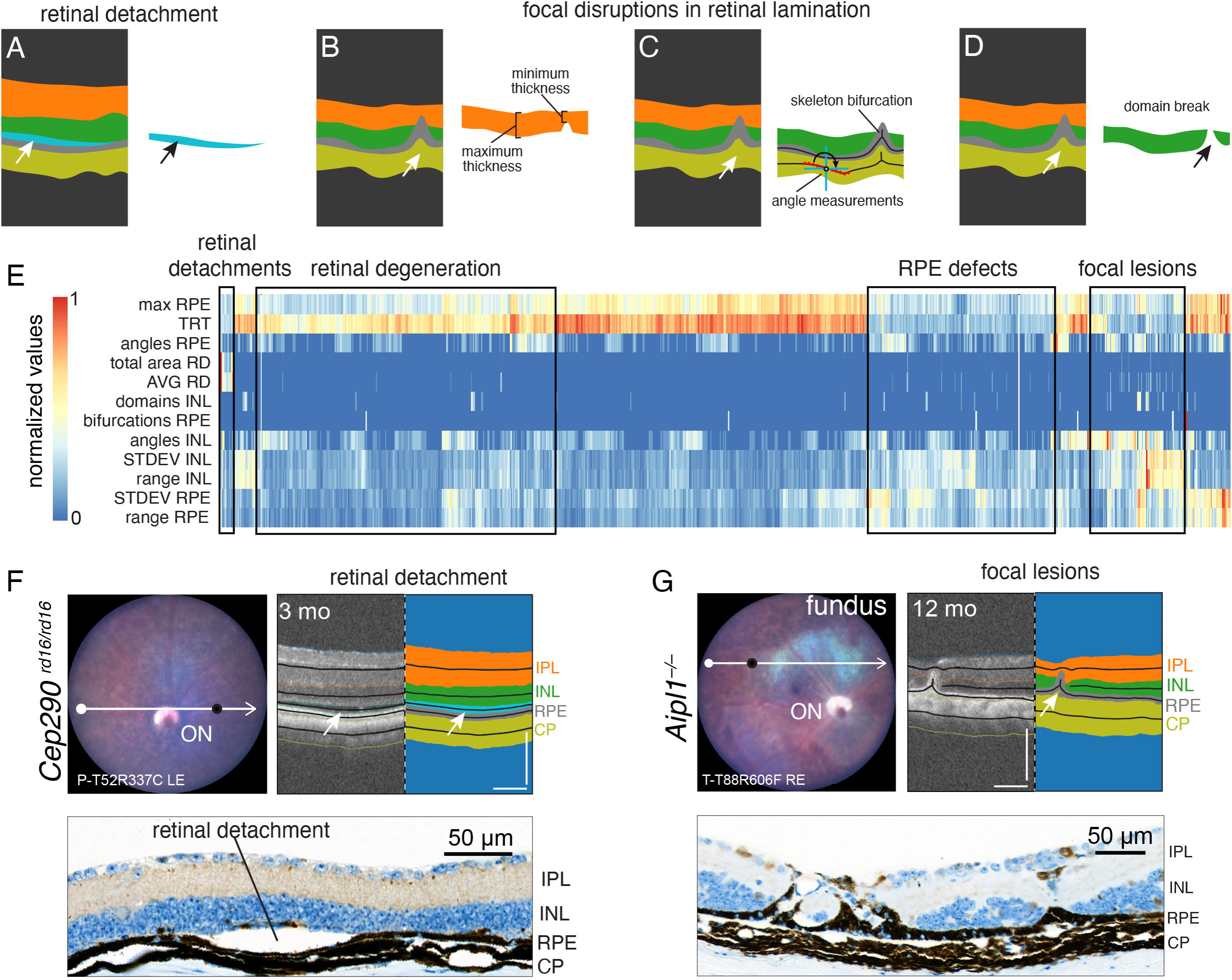
Identification of focal retinal lesions by using C-OCT. A-D) Schematic drawing showing our automated pipeline for segmenting retinal detachments (A), local disruptions in ocular layer thickness (B), acute changes in ocular layer angle and bifurcations of the skeleton of each ocular layer (C), and breaks in the segmentation of individual domains (D). **E)** Heatmap of hierarchical clustered images (n = 1588) of 3-, 6-, 9-, and 12-month-old mice from the following lines: C57BL/6J (n = 18 mice, 237 images), *Rpe65^rd12/rd12^* (n = 15 mice, 200 images) in retinal degeneration cluster, *Ins2^Akita/+^* (female, n = 9 mice, 115 images; male, n = 9 mice, 136 images) only males 6 months and older were included in the retinal degeneration cluster; *Tsc1^lox/lox^*; *Rho*-iCre^Tg^;Tg-*OPN1MW*-Cre^Tg^ (n = 32 mice, 466 images) only images of 12-month-old mice were included in the retinal degeneration cluster; *Cep290^rd16/rd16^* (n = 2 mice, 25 images) were included in the retinal detachment (3-month-old mice only), RPE defects, and focal lesions clusters; and *Aip1^−/−^* (n = 16 mice, 210 images) were included in the RPE defects and focal lesions clusters. **F)** Images of the fundus of a 3-month-old *Cep290^rd16/rd16^* mouse (n = 2/4 mice, 4/8 eyes) with superior scanline (white) and corresponding nasal midline OCT image (location: black dot on fundus) and segmentation showing retinal detachment (arrows; upper panels). Immunohistochemical staining of ARR3 showing retinal detachment in a 3-month-old *Cep290^rd16/rd16^* mouse (lower panel). **G)** Image of a fundus from a 12-month-old *Aipl1^−/−^*(n = 4 mice, 8 eyes) with superior scanline (white) and corresponding superior nasal OCT image (location: black dot on fundus) and segmentation with retinal degeneration (arrow, upper panels). Immunohistochemical staining of CALB showing retinal lesion in a 12-month-old *Aipl1^−/−^*mouse (lower panel). **Abbreviations:** angles, sum of inflections <173°; ARR3, Cone Arrestin 3, AVG, mean thickness across layer; bifurcations, sum of bifurcations per layer; CALB, Calbindin, CP, choroid plexus; domains, sum of domains detected per layer; INL, inner nuclear layer; IPL, inner plexiform layer; max, maximum local thickness; range, range of local thickness; RD, retinal detachment; RPE, retinal pigment epithelium; STDEV, standard deviation of local thickness; total area, the total number of pixels quantified in a layer; TRT, average total retina thickness. OCT images scale bar = 100 μm.

Similarly, the cropped OCT images with local variations in layer thickness, angles, multiple domains, and bifurcations formed a separate cluster that included *Aipl1^−/−^* mice and other strains with rapid retinal degeneration (**Figure 4G**)(35–37). We calculated a sensitivity/specificity score to validate the accuracy of the clusters as they relate to specific retinal phenotypes (**Figure S5D**). Together, these data show the value of integrating multiple measurements from cropped OCT images across the eye to identify different types of focal disruptions in ocular structures in vivo.

### Generalization of the C-OCT Pipeline

To demonstrate the generalized ability of our C-OCT pipeline, we obtained an external dataset of OCT images acquired from C57BL/6J, New Zealand Obese (NZO), and WSB/EiJ mice that are resistant to diet-induced obesity. The time points were 4, 8, 12, and 18 months for each strain with a longitudinal assessment for a sub-cohort of mice within the study. Images were collected on a Phoenix MICRON IV system using the line scan function with 40× averaging. De-identified raw images were run through the C-OCT pipeline. A total of 2136 cropped OCT images passed QC (**Figures 5A**, **S6A-F**). The average individual ocular layer thickness for each mouse showed that the ONL thickness for the NZO mice and WSB/EiJ mice decreased with aging, relative to that of C57BL/6J mice, as reported previously (**Figures 5A,B**, **S6G,H**)(38). The changes in layer thickness of the WSB/EiJ mice were more pronounced than those in NZO mice (**Figure 5A,B**). There was heterogeneity across individual WSB/EiJ mice, and that heterogeneity was exacerbated with aging, tracking longitudinally when sampled (**Figures 5B**, **S6H**). We could not determine whether the variation in layer thickness was regional because only 2 locations were sampled in this external dataset. However, unsupervised hierarchical clustering of features, which was used to identify focal disruptions in ocular structures, highlighted a subset of the WSB/EiJ OCT images (**Figure 5C, Table S4**). Manual review of the cropped OCT images demonstrated that a subset of the WSB/EiJ mice had regions of focal degeneration and RPE infiltration beyond the overall gradual reduction in ONL thickness with age (**Figures 5D,E**, **S6J-L**). Importantly, we did not observe this type of focal retinal degeneration in any of the models used to train and validate our C-OCT pipeline, suggesting that it has broad applicability for diverse preclinical models of retinopathy.

**Figure 5.**
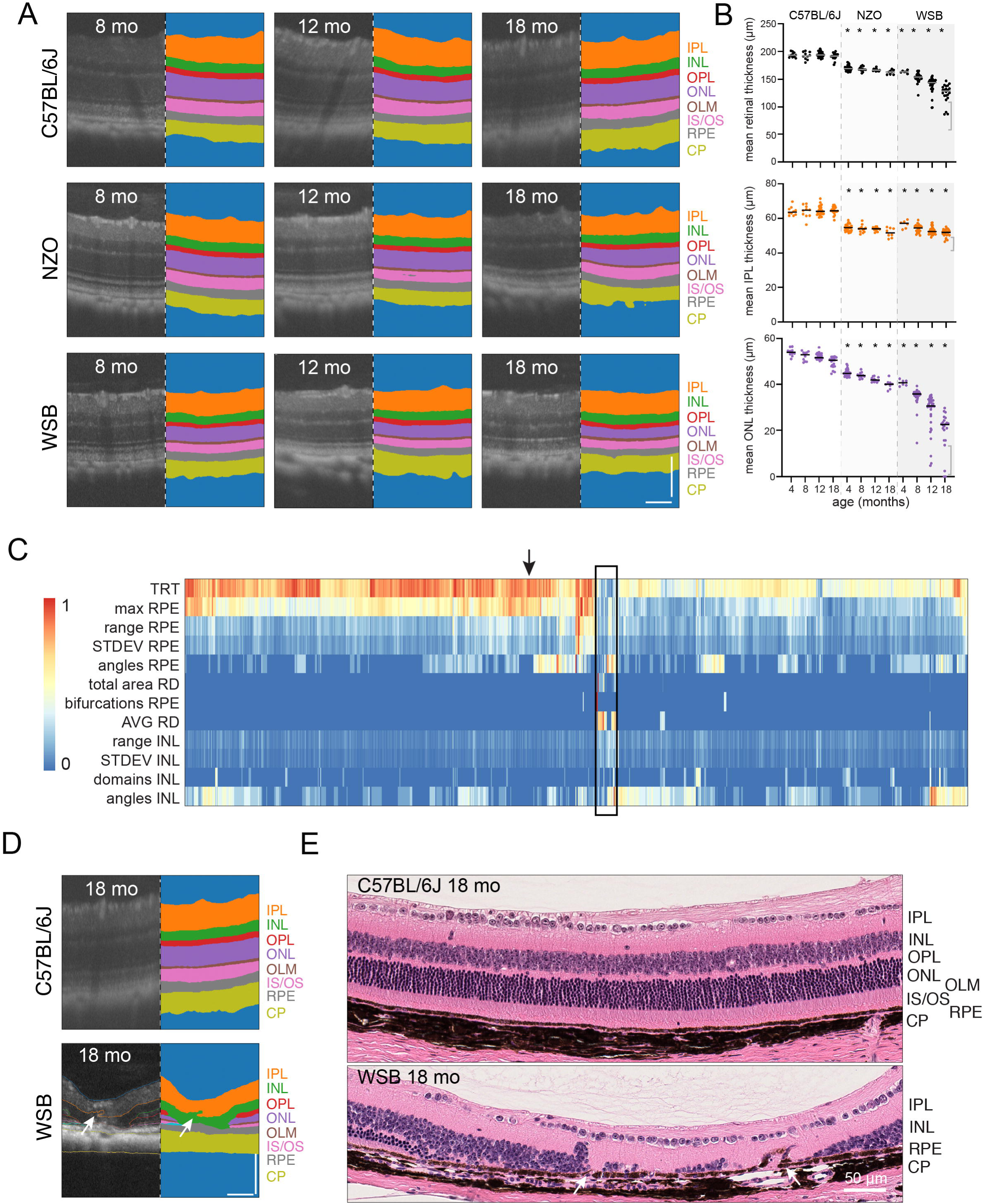
C-OCT is generalizable on independent and diverse datasets. **A)** Representative OCT images of retinal layers with segmentation from 8, -,12-, and 18-month-old C57BL/6J (top), NZO (middle), WSB (bottom) mice. **B)** Quantifications of mean total retinal thickness (top), mean IPL thickness (middle), and mean ONL thickness (bottom) for C57BL/6J (no shading), NZO (light gray shading), and WSB (dark gray shading) mice at 4, 8, 12, and 18 months. Kruskal-Wallis test for significance with FDR adjustment with comparison to C57BL/6J, *p <0.05. **C)** Heatmap of hierarchical clustered images (n = 1061 images) from 12-and 18-month-old mice from the following lines: C57BL/6J (n = 46 mice, 422 images), NZO (n = 19 mice, 168 images) and WSB/EiJ (n = 58 mice, 471 images). The black box notes both RPE defects and focal lesion clusters. **D)** Representative OCT images and segmentation from a C57BL/6J mouse (upper panel) marked with black arrow on heatmap and a WSB mouse with uneven retinal degeneration and a lesion (arrow; lower panel). **E)** Images of H&E-stained sections of 18-month-old C57BL/6J (top) and WSB (bottom) retinae. **Abbreviations:** angles, sum of inflections <173°; AVG, mean thickness across layer; bifurcations, sum of bifurcations per layer; CP, choroid plexus; domains, sum of domains detected per layer; INL, inner nuclear layer; IPL, inner plexiform layer; IS/OS, inner segment/outer segment; max, maximum local thickness; NZO, New Zealand Obese; range, range of local thickness; OLM, outer limiting membrane; ONL, outer nuclear layer; OPL, outer plexiform layer; RD, retinal detachment; RPE, retinal pigment epithelium; STDEV, standard deviation of local thickness; total area, the total number of pixels quantified in a layer; TRT, average total retina thickness; WSB, WSB/EiJ. OCT image scale bar = 100 μm.

## DISCUSSION

OCT is an important diagnostic tool used to track the progression of eye disease in patients and preclinical models. In this study, we developed an end-to-end pipeline called C-OCT for cropping, segmenting, QC filtering, and feature extraction with statistical analysis for murine OCT images. Our pipeline preserves the spatial information of each cropped image, making it ideal for identifying regional and focal lesions associated with retinopathy. C-OCT is computationally efficient, making it possible to analyze thousands of images and extracting millions of features for analysis.

### C-OCT Pipeline

Motion artifacts due to breathing can be a barrier to acquiring high-quality OCT images in mice. Changing parameters in OCT image acquisition such as averaging, taking technical replicates images, duplicate imaging sessions for mice on successive days, or switching the method from a line scan to a volume scan where you acquire up to 512 individual images are ways during experiments to circumvent issues with image quality. In our case, the majority (94.5%) of our OCT images were of sufficient quality for the C-OCT pipeline; therefore, and we employed a de-duplication step to remove replicate crops and avoid a sampling bias in our statistical analysis. We also assessed the C-OCT pipeline segmentation and quantitation of OCT images with 1×, 4×, and 40× averaging on the Phoenix MICRON IV instrument. Despite this, the C-OCT pipeline segmentation and data analyses will be relatively proportional to the quality of the data coming off the instrumentation. Our built-in statistical pipeline generates robust reports so users can properly assess effect size and variation to draw conclusions from their data, regardless of the challenges inherent to working with live subjects. Furthermore, analysis modes can be switched to leverage technical replicates versus individual locations and overall be fine-tuned to the individual’s experimental design.

Comparing the quantitative results from C-OCT to those obtained by ground truth histopathology is also important. Our data on relative thickness of retinal layers were consistent between OCT and histopathologic images, but the absolute values of individual layers differed by as much as 2%. Based on previously published results, we believe our data can be explained by tissue changes associated with formalin fixation and paraffin embedding during preparation of the histologic slides(39, 40).

### Regional Heterogeneity and Focal Lesions

Heterogeneity across the human retina is often measured using the early treatment diabetic retinopathy study grid for location-based macular analyses, peripapillary retinal nerve fiber layer analysis of quadrants around the optic nerve, and finer spatial resolution by using 8×8 grids or clustering algorithms(41–43). Such methods are often employed in AMD progression, glaucoma detection and monitoring, as well as in normal age-related variation(44–47). Although mice lack a macula, they have many examples of regional and focal lesions(48–50). Previous models used either manual location information in their analyses or paired fundus images to register their OCT images(20, 23, 51, 52). Our C-OCT pipeline builds on those prior advances by preserving the scan location and calculating the distance from the optic nerve for each cropped OCT image. The statistical analysis of the features extracted from each eye and aggregated at the mouse level can then be used to identify significant variations that are associated with regional heterogeneity and retinopathy in individual mice, such as those reported here for the *Tsc1^Lox/Lox^;Rho-Cre;OPN1LW-Cre* strain.

Beyond regional variations in retinal layer thickness associated with particular diseases in individual mice over time, a major challenge in the field is identifying focal lesions in individual cropped OCT images. For small numbers of mice, this can be performed manually, but for larger cohorts, automated methods are required. Existing OCT segmentation methods are specialized to identify the focal lesions associated with specific retinopathies(53, 54). To make a more generic tool that uses quantitative feature extraction to identify various focal lesions in very large datasets, we incorporated several approaches to detect focal lesions, including a retinal detachment segmentation mask, analysis of variation in the thickness of retinal layers, skeleton angle measurement, and identification of bifurcations and domain breaks. Data can be analyzed at the image, eye, and mouse level. Using just 12 features, of the hundreds of image features extracted from the data, we identified eyes with focal lesions via unsupervised hierarchical clustering of the cropped OCT images. Importantly, the output from C-OCT can be used for various dimension-reduction-visualization approaches (e.g., PCA, tSNE, UMAP). This is useful not only for identifying focal lesions but also for comparing across diverse models of retinopathy.

### Generalization of C-OCT Output

Our blinded analysis of the previously published OCT data(38) by using the C-OCT pipeline demonstrated the tool’s ability to perform QC, quantitation, and statistical analysis and to identify regional and focal changes associated with various forms of retinopathy. Unlike our OCT imaging at 3 locations across the retina, the external dataset used only 2 locations; this reduced our ability to capture regional heterogeneity. Despite that limitation, we still identified outlier mice and corresponding focal lesions by using hierarchical clustering of image features that were verified with histopathologic analyses. Interestingly, the observed focal lesion cluster included these outlier mice that were longitudinally imaged-supporting that our analysis is viable for longitudinal studies in addition to cross-sectional ones. It will be important to test several instruments and acquisition parameters in future studies to refine and optimize the performance of the C-OCT pipeline and further expand its application to diagnosing other diseases.

### Future Directions

One of the benefits of this approach is its potential for broad application across fields. Here we demonstrate that its efficacy on a variety of mouse models for distinct diseases over stages of degeneration from different institutes. Future studies should focus on application of the pipeline across diverse instrumentation systems. Furthermore, C-OCT is modular and can be broken down where the post-segmentation processing can be cross applied to other segmentation models with minor customization whether it be for mouse, rat, or human. With this modularity, its applicability could extend not just to other species eyes, but perhaps to other tissues with layered structures in general.

Changes in ocular structures are thought to be correlated with nonocular systemic diseases; thus, retinal imaging may detect important biomarkers for diseases that are otherwise difficult to diagnose and monitor. For example, a 29,000-point retinal thickness map across the macula generated in 35,900 patients showed a significant association between retinal thickness and 386 diseases including hyperlipidemia, abnormality of gait, and constipation(9). In rodents, investigators have correlated changes in retinal thickness with Alzheimer disease(55, 56). These and other studies suggest that there are exciting opportunities for developing rigorous pipelines for extracting features from OCT images of preclinical models and patients to identify and validate retinal biomarkers of diverse human diseases.

## METHODS

### Mice and Genotyping

All research was approved by the Institutional Animal Care and Use Committee (IACUC) at St. Jude Childern’s Research Hospital and at The Jackson Laboratory for animals. Mouse models, source information (if they were purchased), and time points of OCT imaging are presented (**Table, S1**). All mice were maintained in the St. Jude Animal Research facility at 71°F, on a 12-hour light/dark cycle, and fed irradiated PicoLab Rodent Diet 20 EXT chow (5R53, LabDiet). Between postnatal days (P) 4 and P7, mice were given a toe identifier (1–100) and genomic DNA was obtained from their tail for genotyping.

Genotyping through PCR and MiSeq analyses were done using direct matches for the respective mutation, and genotypes were designated based on the percentage of mutant reads per total reads detected for genotypes with point mutations. All mice were genotyped for *Pde6b^Rd1^* and *Crb1^Rd8^* upon entrance to the St. Jude colony(57, 58). The *Pde6b^Rd1^*gene was detected in Tg(*OPN1LW*-cre) mice (JAX strain #032911) and was bred out of the line prior to enrolling those mice in the study(33). One to two weeks before OCT imaging, mice were visually inspected for the presence of 2 eyes, eye lesions, blepharitis, rectal prolapse, masses, tumors, and coat color. Mice with eye lesions and/or blepharitis were not used for this study to prevent undue distress on the animal and to eliminate the possibility of contaminating the Phoenix MICRON IV OCT scan head.

The presence of blepharitis was noted only in mice with the Balb/cByJ genetic background, possibly due to increased strain susceptibility(59). Designation of time points and fluctuation around the distinct time point depended on the age of the animal. A maximum of 5 days was tolerated for 3-month-old mice, 2 weeks for 6-month-olds, 3 weeks for 9-month-olds, 3 weeks for 12-month-olds, and 1 month for later time points. All mice in this study had fasting blood glucose and weight recorded for comparison with diabetic mice. All animals housed in at the St. Jude Animal Research Facility participating in this study

### Immunohistochemistry and Histologic Quantifications

Eyes were harvested from 2 male and 2 female mice for the following time points and strains: 3-, 9-, 12-, and 27-month-old C57BL/6J in addition to 3- and 12-month-old *Rpe65^rd12/rd12^* mice, 9-and 12-month-old *Aipl1^−/−^* mice, and 3-month-old *Cep290^rd16/rd16^*. Eyes were then placed immediately in Davidson’s fixative (100 mL 95% alcohol, 200 mL 10% neutral buffered formalin, 100 mL glacial acetic acid, and 300 mL distilled H_2_O) for 3–4 hours. Davidson’s solution rapidly penetrates tissue to ensure proper fixation with fewer artifacts and softens the lens to enable clean sectioning and evaluation. Then eyes were transferred to 10% neutral buffered formalin for 1–2 days. Standard embedding processor settings were then run for infiltration of embedding material and embedding into paraffin blocks. Orientation of the eye was consistent with desired positioning and sectioning with the optic nerve and pupil hole in the ciliary epithelium aligned. Sections were then evaluated by a pathologist to ensure consistency of when and where the start of collected 4-µm consecutive sections could be used for immunohistochemistry or H&E staining. Slides were then scanned at 40× magnification into the HALO Link (Indica Labs, Albuquerque, NM) for visualization and quantitation of layers.

Slides contained 2 consecutive sections for manual quantitation of layer thickness. For each section, 3 regions were assessed, with 3 locations sampled per region for all layer thicknesses. Regions were selected as follows: 1 was near the optic nerve and 2 were symmetrically close to the approximated optical edge, as visualized in SD-OCT. With 18 locations imaged per eye, measurements (in microns) perpendicular to the RPE were then averaged, creating individual values representing layer thickness per mouse for comparisons to C-OCT quantifications. Radar plots for visualization utilized maximum and minimum values derived for all layers and features, across the 15 mouse cohorts, to ensure the resolution of plots were in the context of a wide diversity of phenotypes. RPE infiltrates were counted if they were observed in 3 or more consecutive sections immunostained for CALB2, TH, CALB, and GFAP. While the sections were immunostained RPE infiltrates were not determined by the type of antibody used on each slide. Each infiltrate was tracked across 8 consecutive sections regardless of antibody immunostain in 9- and 12-month-old *Aipl1^−/−^* and C57BL/6J mice. C57BLB/6J, NZO, and WSB eyes were processed as previously described(38). Briefly, mice were anesthetized with tribromoethanol and were transcardially perfused with PBS. Eyes were enucleated and fixed in 37.5% methanol and 12.5% acetic acid in PBS overnight at 4°C. Eyes were paraffin embedded and sectioned. Sections containing the optic nerve head were stained with hematoxylin and eosin and were imaged at 40X magnification.

### Manual SD-OCT Quantitation

Local quantifications across cropped OCT images take ∼415 x-column readings across the skeleton, resulting in 415 readings for each layer thickness. For manual comparison of local quantification across crops, the ROI (region of interest) manager and measurement tool (Fiji Opensource software) were used to acquire ∼200 vertical lengths across x-positions from 3 crops from a single 27-month-old mouse(60). Due to the nonisotropic nature of SD-OCT images, the *x*-length and *y*-length of each ROI were computed with the *x*-length scaled by the 1.62 to isotropic pixels. The hypotenuse was then rederived using the adjusted *x*-length and converted from pixels to microns as the local thickness value. These local thicknesses for each layer were then used to generate the mean, standard deviation, minimum, maximum, interquartile range, and range of the data. Average thicknesses per layer were used to compute the normalized thickness for plotting in radar plots. Minimum and maximums for the radar plots were computed using all data values from the cohort in addition to manual quantification values. One eye per mouse was randomly sampled, and all de-duplicated images associated with that eye were used for quantification. Three representative locations within each cropped OCT image were sampled and measured for thickness of layers by using Fiji software, the conversion methods, and the radar plot generation guidelines mentioned above.

### Statistical Pipeline

To evaluate group-level differences in eye phenotype features, we performed hypothesis testing based on study design and data characteristics. Although statistical pipeline modes for usage of individual eyes or all eyes were included, we found a correlation between features of the left and right eyes. Therefore, average features across eyes for each mouse were used in this manuscript when both eyes were available. Statistical inference was conducted at the mouse. Image- and eye-level features were used for feature extraction, visualization, and descriptive analysis of regional or focal phenotypes, but were not treated as independent units for hypothesis testing. For comparisons between 2 independent groups (e.g., mutant genotype vs. matched control at a fixed time point), we used the Wilcoxon rank-sum test, with effect sizes summarized using Cliff’s delta and false discovery rate (FDR) adjustment applied across features. For comparisons across more than 2 groups (e.g., different time points within a single genotype), we applied the Kruskal-Wallis test, followed by Dunn’s post hoc test with FDR correction for multiple comparisons across timepoints. Effect sizes for Kruskal-Wallis results were reported using epsilon-squared (ε²). These nonparametric tests were used to avoid reliance on normality assumptions. All statistical tests were 2-sided and conducted at a significance level of 0.05. Principal component analysis (PCA) and unsupervised hierarchical clustering were used for exploratory analysis, particularly to visualize the patterns of focal lesions and regional phenotypes. p <0.05.Analyses were performed using R (R Foundation for Statistical Computing, Version 4.2.1). Statistical report folders for each direct comparison were generated with the following items: effect size bar plot of the distribution of features, correlation heatmap of all features, correlation heat map of the top 50 features, summary data table with all statistical findings, PCA plot of eyes, plots for each significant feature between the 2 variables, and an effect size vs –log_10_ (FDR) volcano plot. In addition to each report folder, a sample-counts table for genotypes and time points and a summary table, including all features for each analysis, is provided.

SEM volcano plots utilized the statistical tests outputs log10(P) and log_2_ (fold change of feature SEMs), with a 0.0001 offset added to feature SEMs, between time point–matched C57BL/6J mice and mutants, mice, with the exception of *Abca4^−/−^* mice, which were compared to their respective genetic background Balb/cByJ. A similar comparison was carried out for within-genotype comparisons across time, with each time point compared to the strain’s earliest time point.

### Spectral-Domain Optical Coherence Tomography

While mice underwent initial sedation using isoflurane, Reveal OCT software was started, the white balance correction was carried out, and the reference arm was set to 781. Once mice were sedated, whiskers and eyelashes were trimmed. The following eye drops were then applied to numb the cornea and dilate the pupil for OCT: proparacaine hydrochloride ophthalmic solution, USP 0.5% (24208-730-06, Bausch + Lomb) eye drops, atropine sulfate ophthalmic solution, USP 1% (17478-215-15, AKORN), and tropicamide ophthalmic solution USP 1% (17478-102-12, AKORN). After 5–10 minutes, mice were transferred to the Phoenix MICRON^TM^ IV OCT (40-2273, Phoenix Technologies) platform; sedation was maintained using isoflurane. Dilation eye drops were replaced with GenTeal® lubricant eye gel, severe (4R70, Alcon). and the OCT was centered on the optic nerve by using a transverse scanline. A minimum of 4 OCT images and corresponding fundus images were acquired at the optic nerve, 100 units above, and 100 units below the optic nerve head with 1× or 4× averaging. All left eye and right eye labels recorded in file names were verified using manually recorded time stamps. Each sample was labeled with its experiment number and unique mouse identifier to prevent confusion and enable accurate connection to metadata at a later time. OCT images were then evaluated to determine whether the layers could be discerned by manual review. The images of highest quality were marked “high”; all other images were denoted as “medium,” unless an image was obscured across the full scan, in which case, it was labeled “low” quality and removed from further analysis. Comparisons of C-OCT quantitation of high-quality images with that of medium-quality images were not statistically significant across models, respective time points, or different statistical tests.

OCT scans from Marola *et al.* were acquired using a Phoenix MICRON^TM^ IV OCT with average B-scans taken at 40× averaging (38). Raw images given anonymous strain codes to blind the individual running the analysis and were not reviewed prior to C-OCT. Typically 2 locations were acquired per eye in that dataset, limiting the spatial analysis in C-OCT. Therefore, QC parameters were adjusted to maintain quality eyes with fewer cropped OCT images per eye than the original C-OCT cohort. After completion of the analysis and conclusions, additional blinded individuals graded the fundus and OCT scans for quality, as mentioned above.

### OCT Segmentation

OCT 8-bit TIFFs and paired fundi were fed into the C-OCT pipeline. First, tissue was segmented from background using a model developed using Ilastik(24). Then the scikit-image python package was used with an internal set seed to generate a medial axis(61). Crops were made perpendicular to this medial axis with the center point at the positions of 256, 512, and 768 and dimensions of 256x400 pixels. To annotate individual retinal layers, we developed a custom AI-assisted annotation tool based on MONAI and used it to annotate 296 OCT images, and 219 fundus images. We then trained a Swin-UMamba model using a 5-fold cross-validation schema on this annotated data. **Table S5** summarizes the Dice coefficient for each retinal layer based on this cross-validation. Using the segmented images, we calculated the medial axis for each retinal layer and quantified local thickness along these axes. For fundi analysis, we used our custom AI-assisted annotation tool to annotate the optic nerve head in 219 fundus images. We trained another Swin-UMamba model using 5-fold cross-validation on this data (**Table S5**). To determine the relative location of the OCT crops to the optic nerve, y-coordinates of the scan lines in the fundi images (which denote the location of matching OCT cross-section scans) and the scaled locations of the OCT crop positions.

### Skeletonization and Corresponding Empirical Calculations

The skeleton for the segmented retina was generated using the medial_axis function of the scikit-image python package, with an internal seed set with smoothing(61). Local thickness was calculated in cropped OCT images for each x pixel across a layer by taking the perpendicular distance from the skeleton to the layer boundary and then multiplying it by 2. These values were then used to calculate the average, standard deviation, interquartile range, range, minimum, maximum, and quantiles of layer thickness for the image. Normalized local thicknesses for each ROI, to derive the proportion of each layer, as opposed to the absolute value, were calculated by the average ROI layer thickness divided by either the total retinal thickness or total sclera thickness. Total retina thickness was defined as sum of the IPL, INL, OPL, ONL, ELM, and IS/OS thicknesses, and the total sclera thickness was defined as that of the RPE plus the CHR.

Local angles were calculated by removing the junction points from the skeleton to isolate individual branches. For each pixel on a branch, a straight line was fitted to its neighbors (n = 11) by using a least-squares regression. Pure vertical lines (all dx = 0) were detected and called as 90°, while purely horizontal lines (all dy = 0) were detected and returned as 0°. All other angles were calculated using arctan of the slope of the fitted line. If the least-squares regression failed, an average direction of the nearby pixels (n =11, 5 in each direction) was used to calculate the angle of the center pixel by using the arctan2 of the mean displacements. These returned values generated local angles across each layer’s skeleton and are present in addition to the local thickness for each *x* location, along with *x-* and *y*-coordinates of the skeleton. Inflections along the layer’s skeleton were identified by sign changes in the local sliding window–calculated angle. The location indicated by the sign change was then used to calculate 2 mean angles for the 11 pixels prior to the identified inflection and the 11 pixels after the identified inflection. The angle of inflection was then determined by subtracting the absolute value of the sum of both mean angles from 180°. All vertical angles of 80°-90° were excluded from the analysis to focus on horizontal inflections. Angle bins (<165°, 165°-170°, 170°-173°, 173°-175°, and 175°-177°) were then determined using sampled cropped OCT images with notable inflections. The sensitivity to angle change depends on the layer and may be fine-tuned, depending on the phenotype intended for capture. The number of angles in each bin were then calculated per layer at the image level for further analysis.

### Quality Control Assessment in C-OCT

All cropped OCT images were flagged for QC checking if they contained 2 or more domains in any layer. The number of layers with multiple domains were quantified, and the ratio of the largest domain area versus the total layer area was computed. This value was then averaged across the layers with multiple domains. The images were then split into 2 groups: those with fewer than 8 layers and those with 8 layers. Cropped OCT images with fewer than 8 layers were filtered when they had 2 or more layers with multiple domains and an average largest domain ratio (the largest domain area/the sum of all layer domains areas) less than 75%. Cropped OCT images with 8 or more layers were filtered when either 3 or more layers with multiple domains had an average largest domain ratio less than 98%, when 2 layers with multiple domains had an average ratio of under 85%. Cropped OCT images with 6 or more bifurcations, along the medial axis, across all layers, were filtered. These thresholds were determined by iterative manual assessment using the curated C-OCT cohort.

With the absence of optic nerve midline ROI in the annotation dataset, only 17.2% of optic nerve images passed. As this was not an intended target for analysis, cropped OCT images of optic nerves were excluded from further analysis. Without optic nerve–containing images, the percentage of passing images was 94.84% total and an average of 95.65% across annotated mutants and 92.12% nonannotated mutants. Eyes were filtered if fewer than 2 *y*-anatomical locations were represented, or if fewer than 5 images with OCT segmentation passed QC filtering. To determine if deduplication as opposed to averaging across anatomical regions impacted statistical analyses, the 448 eye level features for C57BL/6J and *Rpe65^rd12/rd12^* at each timepoint were derived both ways and self-compared. While 3897 eye-level features were statistically compared using the Kruskal-Wallis test with adjusted p-value (p <0.05) only the standard deviation of the cumulative total of 4/5 angle bins shown in Supplement SF5A number of inflections in the OLM and the standard deviation of the number of inflections for the widest-angle bin in the choroid. Due to the volume of data, to ensure that our selections of images were truly representative, we did a statistical comparison of extracted features from each individual crop compared to the averaged cohort within an anatomical region for that model and time point. Out of the 3897 calculated features, only 2 were statistical different, thus our representative crops accurately reflect the diverse biology of this large cohort of data.

Our code is provided on github with a downloadable singularity containing the pipeline, alternatively there are also instructions there to build the singularity for it. The post-processing R-script and files used to generate figures with details will also be available on at https://github.com/stjude. Images generated in this study will hosted on the St. Jude website and will be available at 10.5281/zenodo.18623982.

## CONTRIBUTIONS

Danielle Little and Michael Dyer wrote the manuscript. Michael Dyer provided scientific input throughout the project. Danielle Little and Abbas Shirinifard developed the python segmentation algorithms and features for extraction. Abbas Shirinifard wrote the methods portion focused on segmentation. Danielle Little developed the R-post-processing pipeline, acquired, and reviewed OCT images. Marybeth Lupo helped acquire OCT images, bin images, and provided scientific input on R-post processing of OCT images along with review of the manuscript. Chih-Hsuan Wu and Cai Li constructed the statistical pipeline following C-OCT, evaluated all analyses in the paper, and wrote the methods for the statistical pipeline in this manuscript. Danielle Little then integrated their statistical pipeline script into the R-pipeline and converted it into a command line interface for final application. Haoran Chen reviewed both the python segmentation and post-processing along with R post-processing pipelines for quality control and packaged them into a singularity for distribution on github he also reviewed methods. Mellisa Clemons developed manual quantification strategy and wrote methods for quantification of OCT images using Fiji. Michael Maclean, Olivia J. Marola, and Gareth R. Howell provided methods and corresponding data for the histology in **Figure 5E**. Danielle Little processed and analyzed all OCT data in this manuscript.

## ACKNOWLEDGEMENTS

We thank Angie McArthur for editing the manuscript and Ashlley Vallery for her help with optical coherence tomography in this project as a part of animal research services. We also thank Everest Ouyang, Shumei Du, and Sophia Guinocor for helping with mouse enrollment in the study along with Andrea Liao Ramos for organizing images for the associated repository. Funding was provided to M.D. by the NEI (EY037117, EY036689).

## REFERENCES

1. Kiani A, Uyumazturk B, Rajpurkar P, Wang A, Gao R, Jones E, et al. Impact of a deep learning assistant on the histopathologic classification of liver cancer. NPJ Digit Med. 2020;3:23.

2. Lu MY, Chen TY, Williamson DFK, Zhao M, Shady M, Lipkova J, et al. AI-based pathology predicts origins for cancers of unknown primary. Nature. 2021;594(7861):106–10.

3. Pollock LJ, Kitzes J, Beery S, Gaynor KM, Jarzyna MA, Mac Aodha O, et al. Harnessing artificial intelligence to fill global shortfalls in biodiversity knowledge. Nature Reviews Biodiversity. 2025;1(3):166–82.

4. Hu C, Xia Y, Zheng Z, Cao M, Zheng G, Chen S, et al. AI-based large-scale screening of gastric cancer from noncontrast CT imaging. Nat Med. 2025;31(9):3011–9.

5. Ganjgahi H, Haring DA, Aarden P, Graham G, Sun Y, Gardiner S, et al. AI-driven reclassification of multiple sclerosis progression. Nat Med. 2025;31(10):3414–24.

6. Huang W, Shu N. AI-powered integration of multimodal imaging in precision medicine for neuropsychiatric disorders. Cell Rep Med. 2025;6(5):102132.

7. Abramoff MD, Lavin PT, Birch M, Shah N, Folk JC. Pivotal trial of an autonomous AI-based diagnostic system for detection of diabetic retinopathy in primary care offices. NPJ Digit Med. 2018;1:39.

8. Khan Z, Gaidhane AM, Singh M, Ganesan S, Kaur M, Sharma GC, et al. Diagnostic Accuracy of IDX-DR for Detecting Diabetic Retinopathy: A Systematic Review and Meta-Analysis. Am J Ophthalmol. 2025;273:192–204.

9. Jackson VE, Wu Y, Bonelli R, Owen JP, Scott LW, Farashi S, et al. Multi-omic spatial effects on high-resolution AI-derived retinal thickness. Nat Commun. 2025;16(1):1317.

10. Zekavat SM, Jorshery SD, Rauscher FG, Horn K, Sekimitsu S, Koyama S, et al. Phenome-and genome-wide analyses of retinal optical coherence tomography images identify links between ocular and systemic health. Sci Transl Med. 2024;16(731):eadg4517.

11. Sudlow C, Gallacher J, Allen N, Beral V, Burton P, Danesh J, et al. UK biobank: an open access resource for identifying the causes of a wide range of complex diseases of middle and old age. PLoS Med. 2015;12(3):e1001779.

12. Huang D, Swanson EA, Lin CP, Schuman JS, Stinson WG, Chang W, et al. Optical coherence tomography. Science. 1991;254(5035):1178–81.

13. de Boer JF, Cense B, Park BH, Pierce MC, Tearney GJ, Bouma BE. Improved signal-to-noise ratio in spectral-domain compared with time-domain optical coherence tomography. Opt Lett. 2003;28(21):2067–9.

14. Pandya BU, Grinton M, Mandelcorn ED, Felfeli T. RETINAL OPTICAL COHERENCE TOMOGRAPHY IMAGING BIOMARKERS: A Review of the Literature. Retina. 2024;44(3):369–80.

15. Nakazawa M, Hara A, Ishiguro SI. Optical Coherence Tomography of Animal Models of Retinitis Pigmentosa: From Animal Studies to Clinical Applications. Biomed Res Int. 2019;2019:8276140.

16. Zhao D, Lee PY, Wong VHY, Nishimura T, Hoang A, Tran KKN, et al. Retinal Assessment Using In Vivo Electroretinography and Optical Coherence Tomography in Rodent Models of Diabetes. Methods Mol Biol. 2023;2678:37–48.

17. Ahmad NU, Staggers K, Lee K, Mehta N, Domalpally A, Frankfort BJ, et al. Total retinal thickness is an important factor in evaluating diabetic retinal neurodegeneration. BMJ Open Ophthalmol. 2024;9(1).

18. Schneider KJ, Staggers K, Bahrainian M, Channa R. Photoreceptor thickness in UK Biobank participants with and without diabetes mellitus. BMJ Open Ophthalmol. 2025;10(1).

19. Zhang H, Yang B, Li S, Zhang X, Li X, Liu T, et al. Retinal OCT image segmentation with deep learning: A review of advances, datasets, and evaluation metrics. Comput Med Imaging Graph. 2025;123:102539.

20. Ma R, Liu Y, Tao Y, Alawa KA, Shyu ML, Lee RK. Deep Learning-Based Retinal Nerve Fiber Layer Thickness Measurement of Murine Eyes. Transl Vis Sci Technol. 2021;10(8):21.

21. Zeng Y, Zhou J, Li Y, Alvisio B, Czech J, Bissig D, et al. Evaluation of retinal structure changes with AI-based OCT image segmentation for sodium iodate induced retinal degeneration. Front Cell Neurosci. 2025;19:1605639.

22. Antony BJ, Kim BJ, Lang A, Carass A, Prince JL, Zack DJ. Automated segmentation of mouse OCT volumes (ASiMOV): Validation & clinical study of a light damage model. PLoS One. 2017;12(8):e0181059.

23. Sanchez-Rodriguez G, Lou L, Pardue MT, Feola AJ. RetOCTNet: Deep Learning-Based Segmentation of OCT Images Following Retinal Ganglion Cell Injury. Transl Vis Sci Technol. 2025;14(2):4.

24. Berg S, Kutra D, Kroeger T, Straehle CN, Kausler BX, Haubold C, et al. ilastik: interactive machine learning for (bio)image analysis. Nat Methods. 2019;16(12):1226–32.

25. Liu JR, Yang H, Zhou HY, Xi Y, Yu LQ, Li C, et al. Swin-UMamba: Mamba-Based UNet with ImageNet-Based Pretraining. Lect Notes Comput Sc. 2024;15009:615–25.

26. Pang JJ, Chang B, Hawes NL, Hurd RE, Davisson MT, Li J, et al. Retinal degeneration 12 (rd12): a new, spontaneously arising mouse model for human Leber congenital amaurosis (LCA). Mol Vis. 2005;11:152–62.

27. Yoshioka M, Kayo T, Ikeda T, Koizumi A. A novel locus, Mody4, distal to D7Mit189 on chromosome 7 determines early-onset NIDDM in nonobese C57BL/6 (Akita) mutant mice. Diabetes. 1997;46(5):887–94.

28. Gastinger MJ, Kunselman AR, Conboy EE, Bronson SK, Barber AJ. Dendrite remodeling and other abnormalities in the retinal ganglion cells of Ins2 Akita diabetic mice. Invest Ophthalmol Vis Sci. 2008;49(6):2635–42.

29. Han Z, Guo J, Conley SM, Naash MI. Retinal angiogenesis in the Ins2(Akita) mouse model of diabetic retinopathy. Invest Ophthalmol Vis Sci. 2013;54(1):574–84.

30. Barber AJ, Antonetti DA, Kern TS, Reiter CE, Soans RS, Krady JK, et al. The Ins2Akita mouse as a model of early retinal complications in diabetes. Invest Ophthalmol Vis Sci. 2005;46(6):2210–8.

31. Cheng SY, Cipi J, Ma S, Hafler BP, Kanadia RN, Brush RS, et al. Altered photoreceptor metabolism in mouse causes late stage age-related macular degeneration-like pathologies. Proc Natl Acad Sci U S A. 2020;117(23):13094–104.

32. Kwiatkowski DJ, Zhang H, Bandura JL, Heiberger KM, Glogauer M, el-Hashemite N, et al. A mouse model of TSC1 reveals sex-dependent lethality from liver hemangiomas, and up-regulation of p70S6 kinase activity in Tsc1 null cells. Hum Mol Genet. 2002;11(5):525–34.

33. Le YZ, Ash JD, Al-Ubaidi MR, Chen Y, Ma JX, Anderson RE. Targeted expression of Cre recombinase to cone photoreceptors in transgenic mice. Mol Vis. 2004;10:1011–8.

34. Li S, Chen D, Sauve Y, McCandless J, Chen YJ, Chen CK. Rhodopsin-iCre transgenic mouse line for Cre-mediated rod-specific gene targeting. Genesis. 2005;41(2):73–80.

35. Dyer MA, Donovan SL, Zhang J, Gray J, Ortiz A, Tenney R, et al. Retinal degeneration in Aipl1-deficient mice: a new genetic model of Leber congenital amaurosis. Brain Res Mol Brain Res. 2004;132(2):208–20.

36. Chang B, Hawes NL, Hurd RE, Wang J, Howell D, Davisson MT, et al. Mouse models of ocular diseases. Vis Neurosci. 2005;22(5):587–93.

37. Chang B, Khanna H, Hawes N, Jimeno D, He S, Lillo C, et al. In-frame deletion in a novel centrosomal/ciliary protein CEP290/NPHP6 perturbs its interaction with RPGR and results in early-onset retinal degeneration in the rd16 mouse. Hum Mol Genet. 2006;15(11):1847–57.

38. Marola OJ, MacLean M, Cossette TL, Diemler CA, Hewes AA, Reagan AM, et al. Genetic context modulates aging and degeneration in the murine retina. Mol Neurodegener. 2025;20(1):8.

39. Fischer MD, Huber G, Beck SC, Tanimoto N, Muehlfriedel R, Fahl E, et al. Noninvasive, in vivo assessment of mouse retinal structure using optical coherence tomography. PLoS One. 2009;4(10):e7507.

40. Ferguson LR, Grover S, Dominguez JM, 2nd, Balaiya S, Chalam KV. Retinal thickness measurement obtained with spectral domain optical coherence tomography assisted optical biopsy accurately correlates with ex vivo histology. PLoS One. 2014;9(10):e111203.

41. Trinh M, Khou V, Zangerl B, Kalloniatis M, Nivison-Smith L. Modelling normal age-related changes in individual retinal layers using location-specific OCT analysis. Sci Rep. 2021;11(1):558.

42. Photocoagulation for diabetic macular edema. Early Treatment Diabetic Retinopathy Study report number 1. Early Treatment Diabetic Retinopathy Study research group. Arch Ophthalmol. 1985;103(12):1796–806.

43. Paunescu LA, Schuman JS, Price LL, Stark PC, Beaton S, Ishikawa H, et al. Reproducibility of nerve fiber thickness, macular thickness, and optic nerve head measurements using StratusOCT. Invest Ophthalmol Vis Sci. 2004;45(6):1716–24.

44. Grover S, Murthy RK, Brar VS, Chalam KV. Normative data for macular thickness by high-definition spectral-domain optical coherence tomography (spectralis). Am J Ophthalmol. 2009;148(2):266–71.

45. Hwang YH, Kim YY. Glaucoma diagnostic ability of quadrant and clock-hour neuroretinal rim assessment using cirrus HD optical coherence tomography. Invest Ophthalmol Vis Sci. 2012;53(4):2226–34.

46. Shin JW, Uhm KB, Lee WJ, Kim YJ. Diagnostic ability of retinal nerve fiber layer maps to detect localized retinal nerve fiber layer defects. Eye (Lond). 2013;27(9):1022–31.

47. Etheridge T, Liu Z, Nalbandyan M, Cleland S, Blodi BA, Mares JA, et al. Association of Macular Thickness With Age and Age-Related Macular Degeneration in the Carotenoids in Age-Related Eye Disease Study 2 (CAREDS2), An Ancillary Study of the Women’s Health Initiative. Transl Vis Sci Technol. 2021;10(2):39.

48. Barhoum R, Martinez-Navarrete G, Corrochano S, Germain F, Fernandez-Sanchez L, de la Rosa EJ, et al. Functional and structural modifications during retinal degeneration in the rd10 mouse. Neuroscience. 2008;155(3):698–713.

49. Zhao J, Kim HJ, Montenegro D, Dunaief JL, Sparrow JR. Iron overload and chelation modulates bisretinoid levels in the retina. Front Ophthalmol (Lausanne). 2023;3:1305864.

50. Mercau ME, Akalu YT, Mazzoni F, Gyimesi G, Alberto EJ, Kong Y, et al. Inflammation of the retinal pigment epithelium drives early-onset photoreceptor degeneration in Mertk-associated retinitis pigmentosa. Sci Adv. 2023;9(3):eade9459.

51. Inam MG, Inam O, Yang X, Zeng Q, Tezel G. Integrating Retinal Segmentation Metrics with Machine Learning for Predictions from Mouse SD-OCT Scans. Curr Eye Res. 2025;50(5):502–11.

52. Srinivasan VJ, Ko TH, Wojtkowski M, Carvalho M, Clermont A, Bursell SE, et al. Noninvasive volumetric imaging and morphometry of the rodent retina with high-speed, ultrahigh-resolution optical coherence tomography. Invest Ophthalmol Vis Sci. 2006;47(12):5522–8.

53. Karn PK, Abdulla WH. Precision Segmentation of Subretinal Fluids in OCT Using Multiscale Attention-Based U-Net Architecture. Bioengineering (Basel). 2024;11(10).

54. Mellak Y, Achim A, Ward A, Nicholson L, Descombes X. A machine learning framework for the quantification of experimental uveitis in murine OCT. Biomed Opt Express. 2023;14(7):3413–32.

55. Sanchez-Puebla L, de Hoz R, Salobrar-Garcia E, Arias-Vazquez A, Gonzalez-Jimenez M, Ramirez AI, et al. Age-Related Retinal Layer Thickness Changes Measured by OCT in APP(NL-F/NL-F) Mice: Implications for Alzheimer’s Disease. Int J Mol Sci. 2024;25(15).

56. Ferreira H, Martins J, Moreira PI, Ambrosio AF, Castelo-Branco M, Serranho P, et al. Longitudinal normative OCT retinal thickness data for wild-type mice, and characterization of changes in the 3xTg-AD mice model of Alzheimer’s disease. Aging (Albany NY). 2021;13(7):9433–54.

57. Mattapallil MJ, Wawrousek EF, Chan CC, Zhao H, Roychoudhury J, Ferguson TA, et al. The Rd8 mutation of the Crb1 gene is present in vendor lines of C57BL/6N mice and embryonic stem cells, and confounds ocular induced mutant phenotypes. Invest Ophthalmol Vis Sci. 2012;53(6):2921–7.

58. Bowes C, Li T, Frankel WN, Danciger M, Coffin JM, Applebury ML, et al. Localization of a retroviral element within the rd gene coding for the beta subunit of cGMP phosphodiesterase. Proc Natl Acad Sci U S A. 1993;90(7):2955–9.

59. Fukushima A, Yamaguchi T, Ishida W, Fukata K, Taniguchi T, Liu FT, et al. Genetic background determines susceptibility to experimental immune-mediated blepharoconjunctivitis: comparison of Balb/c and C57BL/6 mice. Exp Eye Res. 2006;82(2):210–8.

60. Schindelin J, Arganda-Carreras I, Frise E, Kaynig V, Longair M, Pietzsch T, et al. Fiji: an open-source platform for biological-image analysis. Nat Methods. 2012;9(7):676–82.

61. van der Walt S, Schonberger JL, Nunez-Iglesias J, Boulogne F, Warner JD, Yager N, et al. scikit-image: image processing in Python. Peerj. 2014;2.

