## Supplemental Information and Figures for "Crop-OCT: a Fully Integrated Imageomics Pipeline to Identify Regional and Focal Retinopathy in Murine Models"

### Figure S1

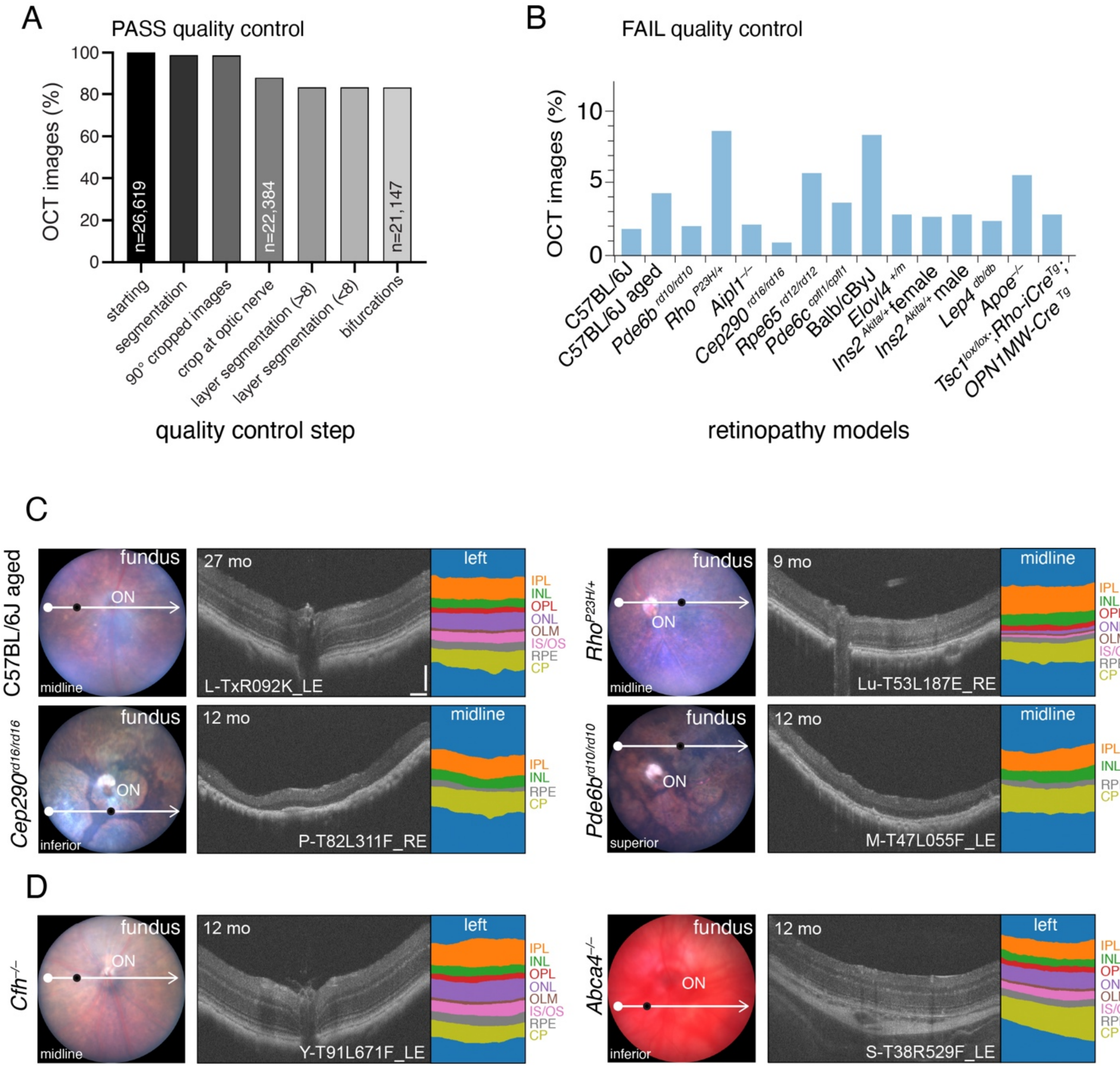
A
PASS quality control
OCT images (%)
0
20
40
60
80
100
n=26,619
n=22,384
n=21,147
starting
segmentation
90° cropped images
crop at optic nerve
layer segmentation (>8)
layer segmentation (<8)
bifurcations
quality control step
B
FAIL quality control
OCT images (%)
0
5
10
C57BL/6J
C57BL/6J aged
Pde6b rd10/rd10
Rho P23H/+
Aipl1 −/−
Cep290 rd16/rd16
Rpe65 rd12/rd12
Pde6c cpfl1/cpfl1
Balb/cByJ
Elovl4 +/m
Ins2 Akita/+ female
Ins2 Akita/+ male
Lep4 db/db
Apoe −/−
Tsc1 lox/lox; Rho-iCre Tg;
OPN1MW-Cre Tg
retinopathy models
C
C57BL/6J aged
fundus
ON
midline
27 mo
L-TxR092K_LE
left
IPL
INL
OPL
ONL
OLM
IS/OS
RPE
CP
Rho P23H/+
fundus
ON
midline
9 mo
Lu-T53L187E_RE
midline
IPL
INL
OPL
ONL
OLM
IS/OS
RPE
CP
Cep290 rd16/rd16
fundus
ON
inferior
12 mo
P-T82L311F_RE
midline
IPL
INL
RPE
CP
Pde6b rd10/rd10
fundus
ON
superior
12 mo
M-T47L055F_LE
midline
IPL
INL
RPE
CP
D
Cfh −/−
fundus
ON
midline
12 mo
Y-T91L671F_LE
left
IPL
INL
OPL
ONL
OLM
IS/OS
RPE
CP
Abca4 −/−
fundus
ON
inferior
12 mo
S-T38R529F_LE
left
IPL
INL
OPL
ONL
OLM
IS/OS
RPE
CP

**Supplemental Figure S1. C-OCT pipeline's segmentation across 15 mouse lines modeling disease. A)** Bar graph depicting the percentage and number of OCT images that passed quality control at each step in the C-OCT pipeline. **B)** Bar graph showing the percentage of OCT images from each mouse line that failed quality control and were removed from further analyses. **C)** Images of fundi with scanline (white) and paired OCT image (location: black dot on fundus) with segmentation from 27-month-old C57BL/6J mice(n = 6 mice, 10 eyes, 552 images; top left panels), 12-month-old *Cep290*$^{rd16/rd16}$ mice (n = 4 mice, 8 eyes, 234 images; bottom left panels), 9-month-old *Rho*$^{P23H/+}$ mice (n = 4 mice, 8 eyes, 260 images; top right panels), 12-month-old *Pde6b*$^{rd10/rd10}$ mice (n = 4 mice, 8 eyes, 287 images; bottom right panels). **D)** Representative mouse lines not included in training annotations are 12-month-old *Cfh*$^{-/-}$ (n = 3 mice, eyes = 6, 227 images) and 12-month-old *Abca4*$^{-/-}$ (n = 7 mice, 14 eyes, 507 images). OCT image scale bar = 100 μm. **Abbreviations:** CP, choroid plexus; INL, inner nuclear layer; IPL, inner plexiform layer; IS/OS, inner segment/outer segment; OLM, outer limiting membrane; ON, optic nerve head; ONL, outer nuclear layer; OPL, outer plexiform layer; RPE, retinal pigment epithelium.

St. Jude - Confidential

### Figure S2

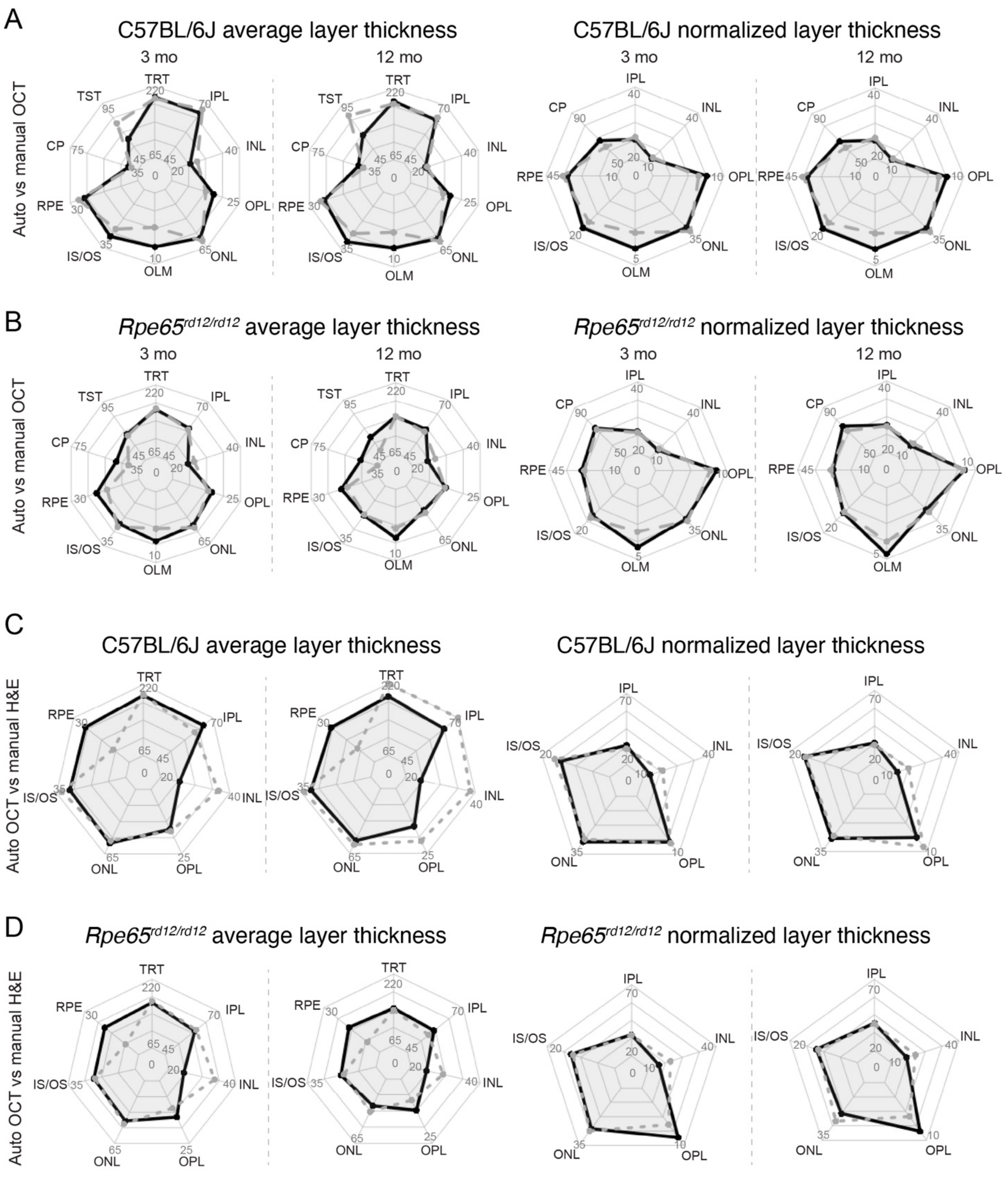
A
C57BL/6J average layer thickness
C57BL/6J normalized layer thickness
3 mo
12 mo
Auto vs manual OCT
TRT
IPL
INL
OPL
ONL
OLM
IS/OS
RPE
CP
TST
B
$Rpe65^{rd12/rd12}$ average layer thickness
$Rpe65^{rd12/rd12}$ normalized layer thickness
C
C57BL/6J average layer thickness
C57BL/6J normalized layer thickness
Auto OCT vs manual H&E
D
$Rpe65^{rd12/rd12}$ average layer thickness
$Rpe65^{rd12/rd12}$ normalized layer thickness

**Supplemental Figure S2. Comparison of C-OCT quantification with manual quantification of OCT images and H&E-stained images**. Radar plots depicting average quantifications of average layer thickness (left panels) and normalized average layer thickness (right panels) from C-OCT quantification (solid black line) and manual quantification (dashed gray line) in 3- and 12-month-old C57BL/6J mice (A) and *Rpe65*$^{rd12/rd12}$ mice (B). C-OCT quantification versus manual H&E quantification (dotted gray line) for C57BL/6J mice (C) and *Rpe65*$^{rd12/rd12}$ mice (D). Minimums and maximums were derived from the C-OCT cohort for each layer.

**Abbreviations:** Auto, C-OCT quantification, CP, choroid plexus; H&E, hematoxylin and eosin; INL, inner nuclear layer; IPL, inner plexiform layer; IS/OS, inner segment/outer segment; OLM, outer limiting membrane; ON, optic nerve head; ONL, outer nuclear layer; OPL, outer plexiform layer; RPE, retinal pigment epithelium; TRT, average total retinal thickness; TST, average total sclera thickness

### Figure S3

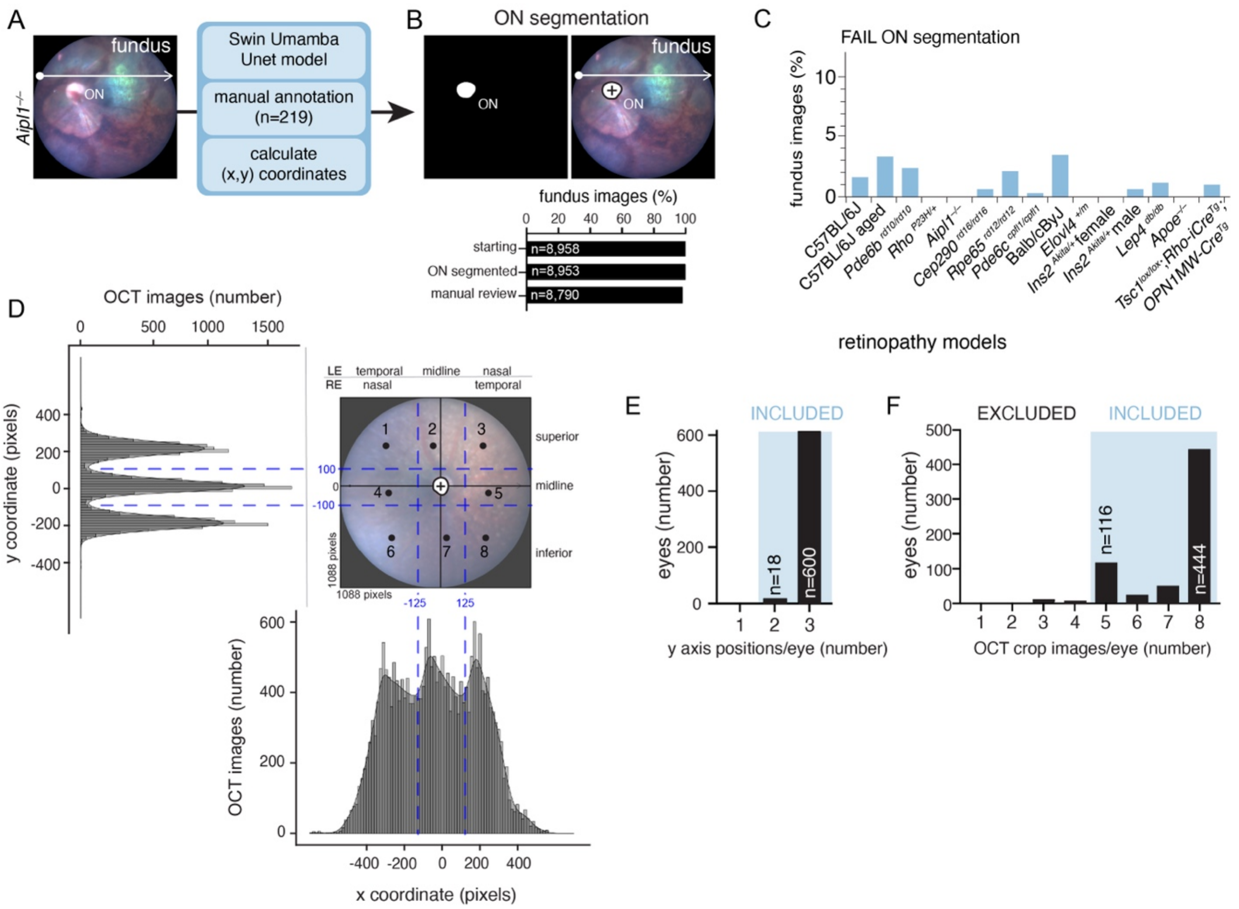

A
fundus
ON
Aipl1−/−
Swin Umamba Unet model
manual annotation (n=219)
calculate (x,y) coordinates
B
ON segmentation
ON
fundus
fundus images (%)
0 20 40 60 80 100
starting n=8,958
ON segmented n=8,953
manual review n=8,790
C
FAIL ON segmentation
fundus images (%)
10
5
0
C57BL/6J
C57BL/6J aged
Pde6b rd10/rd10
Rho P23H/+
Aipl1−/−
Cep290 rd16/rd16
Rpe65 rd12/rd12
Pde6c cpfl1/cpfl1
Balb/cByJ
Elovl4 +/m
Ins2 Akita/+ female
Ins2 Akita/+ male
Lep4 db/db
Apoe−/−
Tsc1 lox/lox;Rho-iCre Tg;
OPN1MW-Cre Tg
retinopathy models
D
OCT images (number)
0 500 1000 1500
y coordinate (pixels)
400 200 0 -200 -400
LE temporal midline nasal
RE nasal temporal
superior
midline
inferior
1 2 3 4 5 6 7 8
100
0
-100
1088 pixels
1088 pixels
-125
125
OCT images (number)
600 400 200 0
-400 -200 0 200 400
x coordinate (pixels)
E
INCLUDED
eyes (number)
600 400 200 0
n=18
n=600
1 2 3
y axis positions/eye (number)
F
EXCLUDED
INCLUDED
eyes (number)
500 400 300 200 100 0
n=116
n=444
1 2 3 4 5 6 7 8
OCT crop images/eye (number)

**Supplemental Figure S3. Maintaining regional complexity in the C-OCT pipeline. A-B)** Representative fundus image input to make the Swin Umamba Unet model (A) and the resulting product of segmentation of fundus optic nerve head (B, upper panel) and quantifications of fundus images maintained with segmentation (B, lower). **C)** Bar plot showing the percentage of images that failed quality control per mouse line. **D)** Density plots showing image distribution from the C-OCT cohort, across a representative fundus image, where the optic nerve head is considered (0,0). The blue dashed lines designate 8 anatomical bins based on their relative distance from the optic nerve head. **E)** Bar plot showing the number of eyes, their *y*-axis anatomical bins represented per eye, and the C-OCT cohort threshold (blue box). **F)** Bar plot showing the number of cropped OCT images per eye, across the C-OCT cohort, and the threshold for an eye to be included for downstream statistical analyses (blue box). **Abbreviation:** ON, optic nerve head

Figure S4

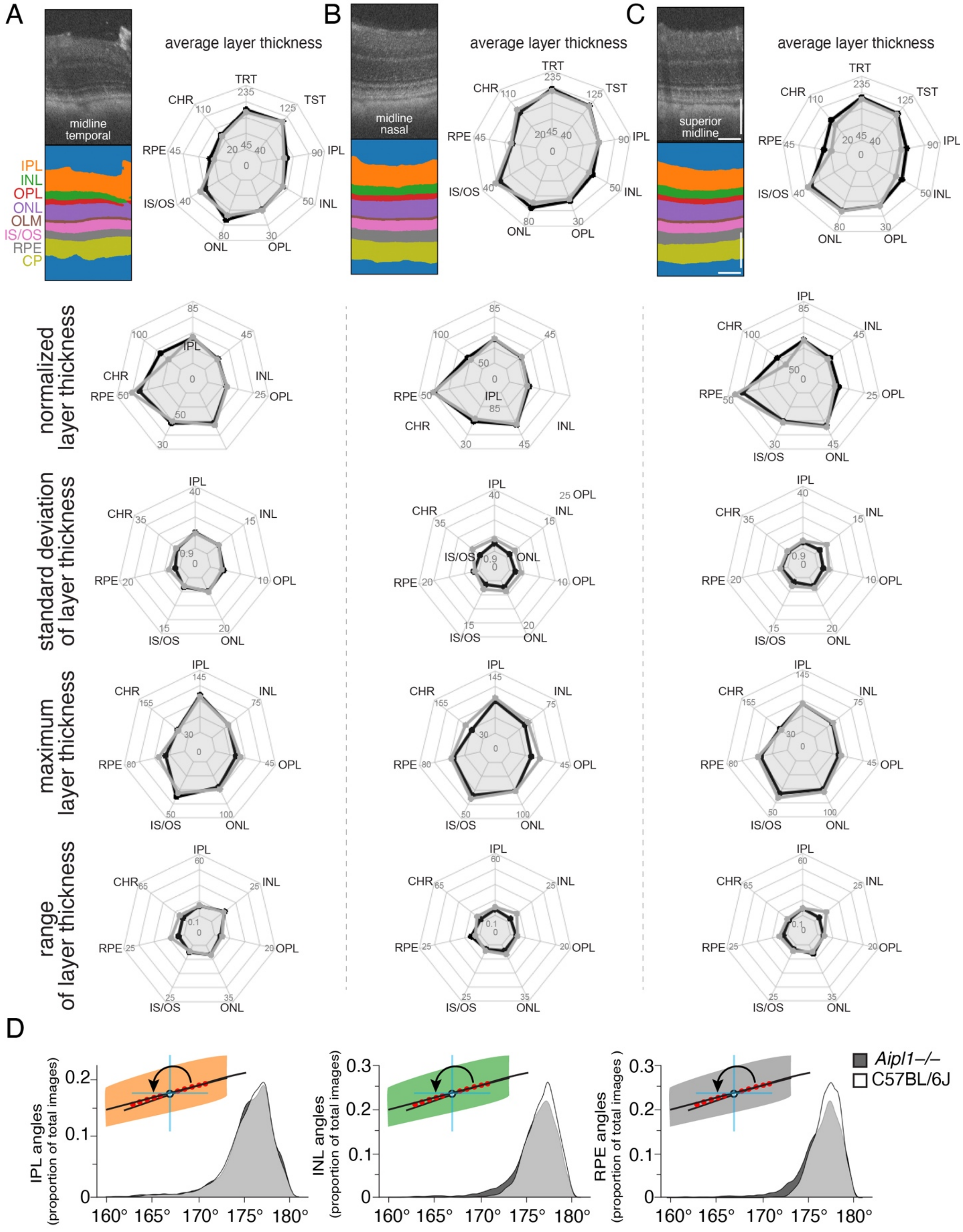
A
B
C
D
midline temporal
midline nasal
superior midline
IPL
INL
OPL
ONL
OLM
IS/OS
RPE
CP
average layer thickness
TRT
TST
IPL
INL
OPL
ONL
IS/OS
RPE
CHR
normalized layer thickness
standard deviation of layer thickness
maximum layer thickness
range of layer thickness
IPL angles (proportion of total images)
INL angles (proportion of total images)
RPE angles (proportion of total images )
160° 165° 170° 175° 180°
Aipl1–/–
C57BL/6J

**Supplemental Figure S4. Manual quantification of local thickness and assessment of local angle inflections for C-OCT quantification. A-C)** Individual OCT image with segmentation and radar plots with C-OCT quantification (black line, n ~ 415 measurements per layer per image) and manual quantification (gray line, n ~200 measurements per layer per image) of local layer thickness from a 27-month-old mouse at the midline temporal position (A), midline nasal position (B), and superior midline position (C). **D)** Density plots of quantifying the angles in degrees of inflections for the IPL (left), INL (middle), and RPE (right) for both *Aipl1*$^{-/-}$ mice (dark gray shading) and C57BL/6J mice (no shading); overlap between the 2 mouse lines is shown in light gray. **Abbreviations:** CP, choroid plexus; H&E, hematoxylin and eosin; INL, inner nuclear layer; IPL, inner plexiform layer; IS/OS, inner segment/outer segment; OLM, outer limiting membrane; ON, optic nerve head; ONL, outer nuclear layer; OPL, outer plexiform layer; RPE, retinal pigment epithelium; TRT, average total retina thickness, TST, average total sclera thickness.

Figure S5

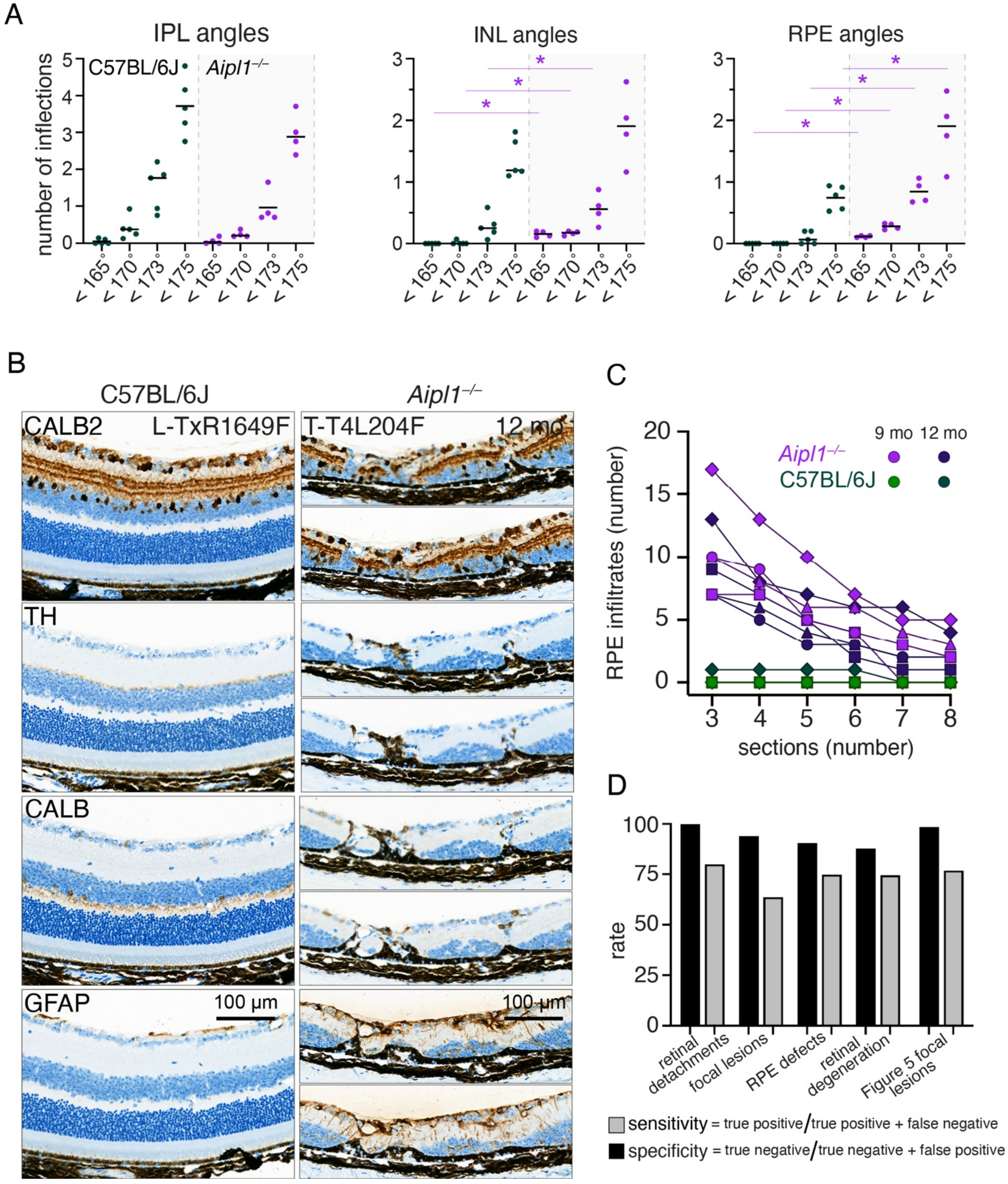
A
IPL angles
INL angles
RPE angles
number of inflections
C57BL/6J
Aipl1−/−
< 165°
< 170°
< 173°
< 175°
B
C57BL/6J
Aipl1−/−
CALB2
L-TxR1649F
T-T4L204F
12 mo
TH
CALB
GFAP
100 µm
C
RPE infiltrates (number)
sections (number)
9 mo
12 mo
Aipl1−/−
C57BL/6J
D
rate
retinal detachments
focal lesions
RPE defects
retinal degeneration
Figure 5 focal lesions
sensitivity = true positive / true positive + false negative
specificity = true negative / true negative + false positive

**Supplemental Figure S5. Detection of focal lesions in *Aipl1*$^{-/-}$ mice. A)** Dot plot of the average number of angles per mouse detected in 9-month-old C57BL/6J mice and *Aipl1*$^{-/-}$ mice (gray) for the IPL (left), INL (center), and RPE (right). Kruskal-Wallis test with adjusted p-values, *p <0.05. **B)** Images of immunohistochemical staining of the RPE across multiple consecutive sections from 12-month-old C57BL/6J mice (left, n = 4 mice, 8 images) and *Aipl1*$^{-/-}$mice (right, n = 4 mice, 8 images). **C)** Corresponding quantification of RPE lesions and the number of consecutive sections on which they were detected in 9- and 12-month-old C57BL/6J mice (both strains and ages: n = 4 mice, 8 images). **D)** Bar plot depicting calculated sensitivity and specificity for visually defined clusters observed in Figure 4E and Figure 5C. the number of images from each genotype and time point for the 3 boxed regions in Figure 3B. **Abbreviations:** CALB2, calretinin; CALB, calbindin 1; GFAP, glial fibrillary acidic protein; INL, inner nuclear layer; IPL, inner plexiform layer; RPE, retinal pigment epithelium; TH, tyrosine hydroxylase

Figure S6

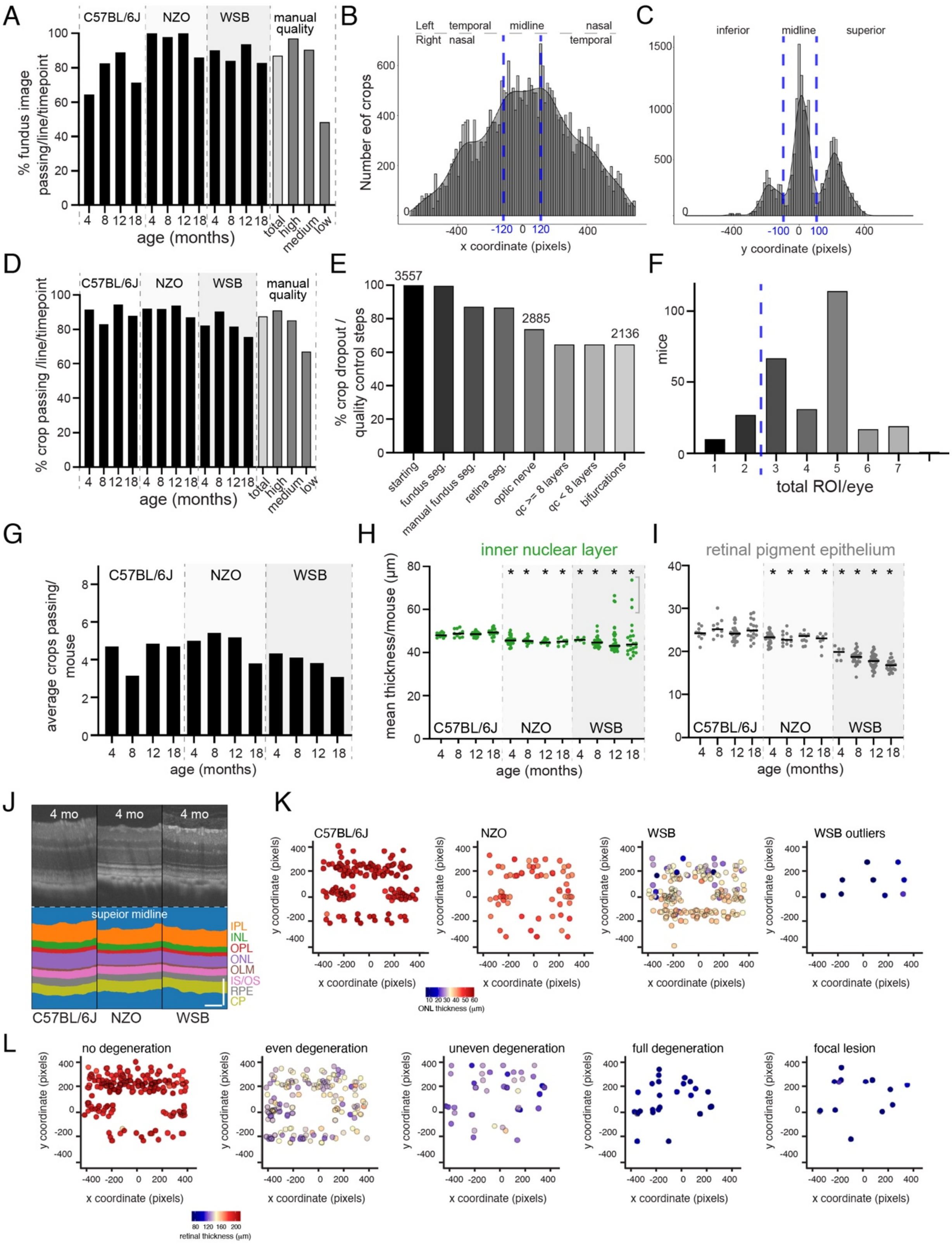

A
C57BL/6J
NZO
WSB
manual quality
% fundus image passing/line/timepoint
age (months)
B
Left
Right
temporal
nasal
midline
nasal
temporal
Number eof crops
x coordinate (pixels)
C
inferior
midline
superior
y coordinate (pixels)
D
% crop passing /line/timepoint
age (months)
E
% crop dropout / quality control steps
3557
2885
2136
starting
fundus seg.
manual fundus seg.
retina seg.
optic nerve
qc >= 8 layers
qc < 8 layers
bifurcations
F
mice
total ROI/eye
G
average crops passing/ mouse
age (months)
H
inner nuclear layer
mean thickness/mouse (µm)
age (months)
I
retinal pigment epithelium
age (months)
J
4 mo
superior midline
IPL
INL
OPL
ONL
OLM
IS/OS
RPE
CP
K
WSB outliers
y coordinate (pixels)
x coordinate (pixels)
ONL thickness (µm)
L
no degeneration
even degeneration
uneven degeneration
full degeneration
focal lesion
retinal thickness (µm)

**Supplemental Figure S6. C-OCT pipeline execution, quality control metrics, and quantification of an external validation dataset. A)** Bar graph depicting the percentage of segmented images of the Fundus optic nerve head that passed quality control (QC) review per time point per line for C57BL/6J, NZO (light gray), and WSB/EiJ (gray) mice, and in the context of manual QC review calling of high, medium, or low quality. **B)** Histogram depicting the distribution of OCT images across the y-axis, and where they fall in reference to the C-OCT pipeline's anatomical bins (dark blue dashed lines). **D)** Bar graph depicting OCT-segmented images that passed QC review per time point per line for C57BL/6J, NZO (light gray), WSB/EiJ (gray) mice, and in the context of manual QC review calling of high, medium, or low quality. **E)** Bar graph of percentage and number of cropped OCT images filtered at each QC step. **F)** Bar graph showing the number of mice with images per eye across the dataset, with a threshold of 3 (blue dashed line). **G)** Bar plot of the average number of images that passed QC review per mouse for C57BL/6J, NZO, and WSB/EiJ mice across time. **H-I)** Dot plot of the mean layer thickness per mouse (each dot) for 4-, 8-, 12-, and 18-month-old C57BL/6J (n = 10, 10, 27, 19 mice per time point) NZO (n = 32, 12, 11, and 8 mice per time point), and WSB/EiJ mice (n = 6, 25, 36, 22 mice per time point) for the INL (H) and RPE (I). **J)** OCT images and respective segmentation for 4-month-old C57BL/6J, NZO, and WSB/EiJ mice (scale bar = 100 μm). **K)** Eye maps of ONL thickness plots of all de-duplicated images (points) from each eye on the optic nerve coordinate grid, with the mean ONL thickness represented by the hue of each point for all 12-month-old C57BL/6J (n = 19, left), NZO (n = 8, center) and WSB/EiJ (n = 22, center right) mice and 12-month-old WSB outliers (n = 3, far right). **L)** Eye maps of retinal thickness plots of images (dot), with the mean total retinal thickness represented by the hue of each dot within manual graded bins of no degeneration (n = 471 images; far left), relatively even amounts of degeneration across the layers (n = 377 images; center left), and uneven degeneration with large variance and a consistent loss of thickness across the layer (n = 64 images; center). Retinal thickness plots of images with full degeneration, where outer layers were not observed (n = 26 images; center right) and those with RPE infiltration into the retina (n = 14 images; far right) are also shown. **Abbreviations:** CP, choroid plexus; INL, inner nuclear layer; IPL, inner plexiform layer; IS/OS, inner segment/outer segment; NZO, New Zealand Obese; ONL, outer nuclear layer; OPL, outer plexiform layer; ROI, region of interest; RPE, retinal pigment epithelium; retinal thickness, average total retina thickness; WSB, WSB/EiJ.

St. Jude - Confidential
